# Ecological context shapes the interpretation of biodiversity intactness metrics

**DOI:** 10.64898/2026.10.26.761996

**Authors:** Thiago Gonçalves-Souza, Reginaldo Gusmão, Dimitris Gounaridis, Elnaz Amirahmadi, Dunya Diyadawagamage, Geoffrey M. Lewis, Joshua Newell, Peter B. Reich, Dominick Stoops, Kai Zhu, Benjamin P. Goldstein

## Abstract

Developing biodiversity metrics that are both operational for decision-making and ecologically meaningful remains challenging. Mean Species Abundance (MSA) is widely used to quantify biodiversity intactness in applications including biodiversity scenario modeling, spatial assessment, policy and planning, and life cycle assessment (LCA). MSA compares species abundance under disturbed conditions with their abundance under a reference condition, but the pressure–impact relationships used to calculate it are generalized across studies and may not adequately represent variation among ecological contexts or taxonomic groups. This is particularly relevant when MSA is transferred across regions or used to support comparisons and environmental decision-making. We compiled biodiversity data from 7,064 forest fragments containing 21,761 species of arthropods, amphibians, birds, mammals, squamates, and plants across 11 biomes worldwide. We evaluated how MSA relates to broader ecological outcomes, including local and regional species richness and ecosystem productivity, and whether these relationships vary across ecological contexts (vegetation condition, landscape structure, and biome) and taxonomic groups. MSA was positively associated with local species richness and ecosystem productivity across all sites, but these associations weakened, disappeared, or reversed depending on ecological context and taxonomic group. For example, the MSA–productivity association held in only five of 11 biomes, and the MSA–regional richness association was negative in temperate conifer and boreal forests but positive in four other biomes. We propose a simple decision guide linking assessment goals, contextual evaluation, and interpretation to help identify when MSA can be used directly, when ecological context should guide its interpretation, and when complementary ecological information is needed. These findings may extend to other biodiversity metrics that compare impacted conditions with reference states and apply generalized impact relationships across ecological contexts and taxonomic groups.

## 1. Introduction

Resource extraction is reshaping biodiversity and ecosystems globally, threatening around one million species with extinction (IPBES 2019, Jaureguiberry *et al* 2022, Pereira *et al* 2024). These impacts affect human well-being through significant deterioration in ecosystem services, such as water and food provisioning (Díaz *et al* 2006, Cardinale *et al* 2012, Naeem *et al* 2016). There is an urgent need for indicators that can inform environmental decision-making (Burgess *et al* 2024) and mitigate losses to biodiversity and ecosystem services (Feld *et al* 2009, Stevenson *et al* 2021).

Life cycle assessment (LCA) has emerged as the de facto decision support tool to quantify the environmental and biodiversity impacts of production and consumption (Sala *et al* 2020, Hellweg *et al* 2023, Jegen 2024). Yet how well do biodiversity indicators used in LCA reflect real-world ecological outcomes? Mean Species Abundance (MSA) is used in GLOBIO, a global biodiversity model for policy support, to quantify biodiversity intactness across applications including scenario analysis, policy and planning, and spatial biodiversity assessment (Alkemade *et al* 2009, Schipper *et al* 2020). MSA is also incorporated into some LCA biodiversity assessment approaches through characterization factors (Schipper *et al* 2020). Despite its broad application, MSA estimates are rarely validated against empirical field data (Lüscher *et al* 2017, Hayashi 2020, VanderWilde and Newell 2021). Moreover, the characterization factors that convert biodiversity stressors (e.g., land use conversion) in these models often rely on globally or nationally averaged values that do not capture the spatiotemporal variation in land use and biodiversity responses (VanderWilde and Newell 2021, Avila-Ortega *et al* 2025). Consequently, whether these proxies accurately reflect biodiversity responses across ecological contexts and taxonomic groups remains unclear (Damiani *et al* 2023). This validation gap is even greater for ecological outcomes beyond species richness, including ecosystem processes (Teixeira *et al* 2016, Weinzettel *et al* 2019, Song *et al* 2025). Importantly, this gap persists despite the growing availability of high-resolution global and regional biodiversity datasets (Kissling *et al* 2018, Wüest *et al* 2020, Clements *et al* 2024, Dasgupta *et al* 2026, El-Gabbas 2026).

This challenge partially stems from differences in how biodiversity is conceptualized and measured in ecology and model-based biodiversity assessment (Damiani *et al* 2023). Ecologists use metrics of species richness, abundance, and composition to characterize the state of ecological communities at fine spatial scales and specific locations (Santini *et al* 2017, Avila-Ortega *et al* 2025). In contrast, biodiversity indicators used in frameworks such as GLOBIO and LCA simplify ecological responses to support broader environmental assessment and decision-making, including estimates of the biodiversity impacts of production and consumption. (Damiani *et al* 2023, Avila-Ortega *et al* 2025). Within LCA, ongoing initiatives such as BAMBOO and RAINFOREST are broadening biodiversity assessment by incorporating additional biodiversity dimensions and more spatially explicit approaches (Dürr *et al* 2026, Zhang *et al* 2026). These conceptual and spatial mismatches nevertheless limit the ability of model-based biodiversity indicators to capture actual ecological responses across different ecological contexts.

MSA, which is often used directly or incorporated into LCA indicators, compares species abundance under disturbed conditions (e.g., agriculture) relative to pristine conditions, for grid cells or other spatial units (e.g., 1,000 ha of converted forest) (Alkemade *et al* 2009, Schipper *et al* 2020). By comparing changes in abundance across multiple species within a spatial unit, MSA represents an average response. The calculation and interpretation of this response, however, rely on two important assumptions. The first assumption is that reference species are undisturbed by human activities in pristine or primary vegetation (Alkemade *et al* 2009, Schipper *et al* 2020). Second, when MSA is projected globally, equivalent pressure-response relationships are applied with the same weight across spatial units, irrespective of differences in climate, biome, or species composition (Alkemade *et al* 2009, McNellie *et al* 2020). Furthermore, defining an undisturbed reference state is challenging because deviations from reference conditions can vary across land-use and climatic contexts (McNellie *et al* 2020). Since ecological communities can respond differently under contrasting land-use and climatic conditions (Etard and Newbold 2024, Bevan *et al* 2025), the ability of MSA to reflect biodiversity responses may also vary across these ecological contexts (Arets *et al* 2014).

Although MSA is sometimes interpreted in the context of ecosystem integrity (Marques *et al* 2021, Kuipers *et al* 2025, Sidemo-Holm *et al* 2026), it directly measures biodiversity intactness rather than ecosystem functioning. Ecosystem functions, such as carbon storage or productivity, represent a distinct dimension of ecosystem condition and are often assessed alongside biodiversity indicators (Hillebrand *et al* 2018, Dıaz and Cabido 2001, Feld *et al* 2009). This distinction is important because MSA captures average species abundances, whereas ecosystem functions often depend on the abundance of dominant species or key functional groups, and on functional complementarity among species (Reich *et al* 2012, Winfree *et al* 2015, Zhang *et al* 2025). Clarifying the extent to which biodiversity intactness measured by MSA corresponds to ecosystem functioning is therefore important for interpreting MSA in broader ecological assessment and LCA.

In this study, we investigate the contexts in which MSA reflects different ecological outcomes in forests. Specifically, we address the following questions: (i) How closely does MSA reflect local and regional species richness and ecosystem productivity functioning forests? (ii) How do these relationships vary across ecological contexts (vegetation properties, landscape structure, and biome), taxonomic groups, and spatial scales? To do this, we test alignment between MSA and empirical data on local species richness, regional species richness, and forest productivity from 7,064 forest fragments containing 21,761 species across six taxa in 11 biomes worldwide. By answering these questions, we provide an empirical evaluation of whether relationships between MSA and other ecological outcomes are transferable across systems or conditional on the contexts in which MSA is interpreted. These findings have direct implications for how MSA-based biodiversity indicators are interpreted and applied in LCA.

## 2. Methods

We used complementary species- and ecosystem-targeted approaches to evaluate how well MSA reflects biodiversity and ecosystem functioning across different ecological outcomes. The fundamental sampling unit was a continuous forest (1,000 - 300,000 ha) or forest fragment (<800 ha) (Gonçalves-Souza *et al* 2025a, 2025b). We used the geographic centroid of each forest as the sampling location from which to extract MSA, local and regional species richness, biome, ecosystem productivity, remotely-sensed vegetation indices, and landscape metrics (Fig. 1).

**Figure 1.**
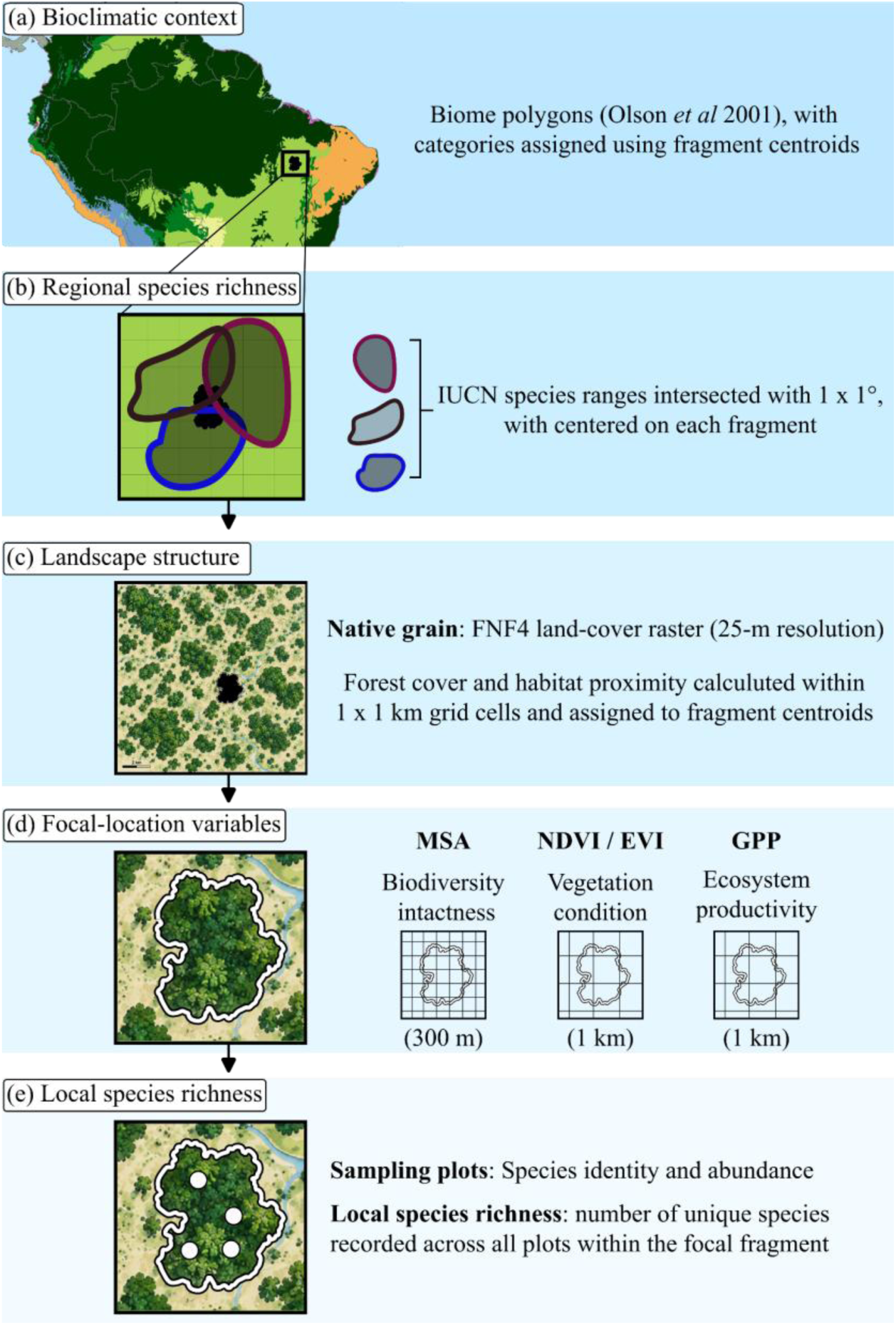
Spatial scales and resolutions of biodiversity and environmental data used in the study. The schematic illustrates how ecological information from different spatial resolutions and geographic extents was integrated using forest-fragment centroids as reference locations. From broad to local scales, the figure shows (a) biome classification representing bioclimatic context; (b) regional species richness derived from IUCN species ranges; (c) landscape structure characterized by forest cover and habitat proximity; (d) model-derived MSA, vegetation indices (NDVI and EVI), and cumulative DHI-GPP; and (e) local species richness from field observations.

### 2.1. Mean species abundance (MSA)

The first step in calculating MSA is to divide, for each species occurring in the reference assemblage, its abundance under the disturbed condition by its abundance under the corresponding reference condition, with values greater than 1 truncated at 1. MSA is then computed as the mean of these values across all species (Alkemade et al 2009, Schipper et al 2020). Species present in the reference species condition that are absent from the disturbed community contribute zero, whereas species occurring only in the disturbed community are excluded. Consequently, MSA ranges from 0 to 1, where 1 indicates that every reference species maintains or exceeds its reference abundance, and 0 indicates the absence of all reference species in the disturbed condition (Schipper et al 2020). The GLOBIO model provides a global MSA map as a GeoTIFF raster at a spatial resolution of 10 arc-seconds, approximately 300 × 300 m at the equator (Schipper et al 2020). We used the 2015 MSA raster and extracted the corresponding modeled MSA value for each sampling location. Full information about the combination of pressure-response relationships to calculate MSA are provided in the Supplementary Material.

### 2.2 Species-targeted approach

#### 2.2.1 Local species richness

We combined site-level biodiversity data for species richness and abundance at each sampled forest fragment from two global databases (figure S1), LandFrag (Gonçalves-Souza *et al* 2025b) and PREDICTS (Hudson *et al* 2014). Taxonomic groups comprised vascular plants, arthropods, birds, mammals, amphibians, and squamates. We included georeferenced community-level studies conducted in continuous forest or forest fragments for which species abundance and composition data were available. We retained 108 studies from LandFrag and 193 studies from PREDICTS that met these criteria. The combined dataset contained 7,064 sampling locations from LandFrag (n = 1,726 sampling locations and 8,278 species) and PREDICTS (n = 5,338 sampling locations and 13,707 species). We calculated local species richness (number of species recorded at each location), total number of individuals collected on each location, and average abundance across species. We included these variables to partially account for differences in sampling effort among studies (Supplementary Material), while recognizing that differences in sampling design could not be fully standardized across datasets (Chase *et al* 2020, Gonçalves-Souza *et al* 2025a).

#### 2.2.2 Regional species richness

We used geographical range polygons provided by the International Union for Conservation of Nature (IUCN 2026) to estimate regional species richness. We counted the number of unique species whose geographic range polygons intersected with a 1° × 1° spatial window. Range maps were assembled for plants, mammals, birds, amphibians, and squamates and transformed to a common coordinate reference system before spatial intersection. Since these polygons represent mapped geographic distributions rather than species directly observed at the sampling location, we refer to this response as regional species richness. Unlike local species richness, regional richness represents biodiversity over a substantially broader spatial extent than the MSA estimates. Consequently, analyses of regional species richness evaluated among-study relationships with MSA across different spatial scales rather than within-study relationships at comparable spatial extents (figure 1).

### 2.3 Ecosystem-targeted approach

We used the cumulative Dynamic Habitat Index derived from MODIS gross primary productivity (DHI-GPP) at 1-km resolution (Radeloff *et al* 2019, Liu *et al* 2026). GPP represents the total carbon fixed by primary producers through photosynthesis before accounting for autotrophic respiration (Regos *et al* 2022). The cumulative DHI-GPP summarizes GPP across the annual phenological cycle and was used here as an indicator of ecosystem photosynthetic productivity at each sampling location (Bao *et al* 2022, Beer *et al* 2010). Hereafter, we refer to cumulative DHI-GPP as ecosystem productivity. More details on its calculation are provided in the Supplementary Material (see also figure 1).

### 2.4 Defining ecological context

We evaluated whether the relationships of MSA with species richness and ecosystem productivity varied across three dimensions of ecological context: remotely sensed vegetation properties, landscape structure (forest cover and habitat proximity), and bioclimatic context (biome) (figure 1, table S1).

#### 2.4.1. Vegetation condition

We used the Normalized Difference Vegetation Index (NDVI) and Enhanced Vegetation Index (EVI) as remotely sensed measures of vegetation properties (figure 1). These indices provide complementary information on vegetation greenness, photosynthetic activity, and canopy characteristics. We used the cumulative Dynamic Habitat Index components derived from NDVI and EVI between 2013 and 2022 extracted at 1-km resolution to represent longer-term vegetation conditions surrounding each sampling location (Radeloff *et al* 2019).

#### 2.4.2. Landscape structure

We characterized landscape structure using forest cover and habitat proximity (figure 1). We used the 2020 JAXA ALOS PALSAR/PALSAR-2 Forest/Non-Forest (Japan Aerospace Exploration Agency (FNF4) 2016) dataset at 25m spatial resolution, combining dense and non-dense forest classes as forest. Following Gounaridis et al. (2020), we divided the global forest layer into 1 × 1 km grid cells and treated each cell as an independent landscape in FRAGSTATS. Within each cell, forest cover was calculated as the percentage of the landscape occupied by forest, while habitat proximity was quantified using the mean proximity index, which incorporates the size and distance of neighboring forest patches (McGarigal *et al* 2023, Neel *et al* 2004).

#### 2.4.3. Bioclimatic context

We used the global map of terrestrial ecoregions defined by Olson et al. (2001), which classifies terrestrial ecosystems into 14 biomes, but retained only the 11 biomes with sufficient data for analysis. Since these biomes represent broad differences in climatic and vegetation conditions, we used the biomes as a fixed effect in the models to evaluate whether the relationships between MSA and the ecological outcomes varied among biomes (figure 1). Hereafter, we collectively refer to vegetation condition, landscape structure, and bioclimatic context as ecological context. Likewise, we refer to local and regional species richness and ecosystem productivity as ecological outcomes.

### 2.5 Data analysis

We evaluated how closely MSA was associated with local and regional species richness and ecosystem productivity by fitting generalized linear mixed-effects models using the R package glmmTMB (Brooks *et al* 2017). Species richness was modeled as count data using a negative binomial distribution with the NB2 parameterization and a log link to accommodate overdispersion (Hardin and Hilbe 2018), while ecosystem productivity was log-transformed and modeled using a Gaussian error distribution.

To account for ecological context, we fitted interaction models between MSA and each ecological-context variable. MSA and continuous ecological-context variables were standardized before analysis. Study identity was included as a random intercept to account for non-independence among observations from the same study. The taxonomic group was included as a fixed effect in models of species richness. Full model equations and sensitivity analyses are provided in the Supplementary Material.

## 3. Results

We found that higher modeled MSA, representing greater biodiversity intactness, was positively associated with local species richness (figure 2A) and ecosystem productivity (figure 2C), but negatively associated with absolute regional species richness (figure 2B, table S2). Accounting for ecological context substantially altered the magnitude and, in some cases, the direction of these relationships. For local species richness, positive associations were concentrated mainly in tropical biomes, especially tropical and subtropical moist broadleaf forests (except for birds) and tropical and subtropical dry broadleaf forests (only for arthropods) (figure 3), as well as for birds and mammals in tropical and subtropical grasslands, savannas, and shrublands. In the remaining biome–taxon combinations, there was no clear relationship between MSA and local species richness (figure 3).

**Figure 2.**
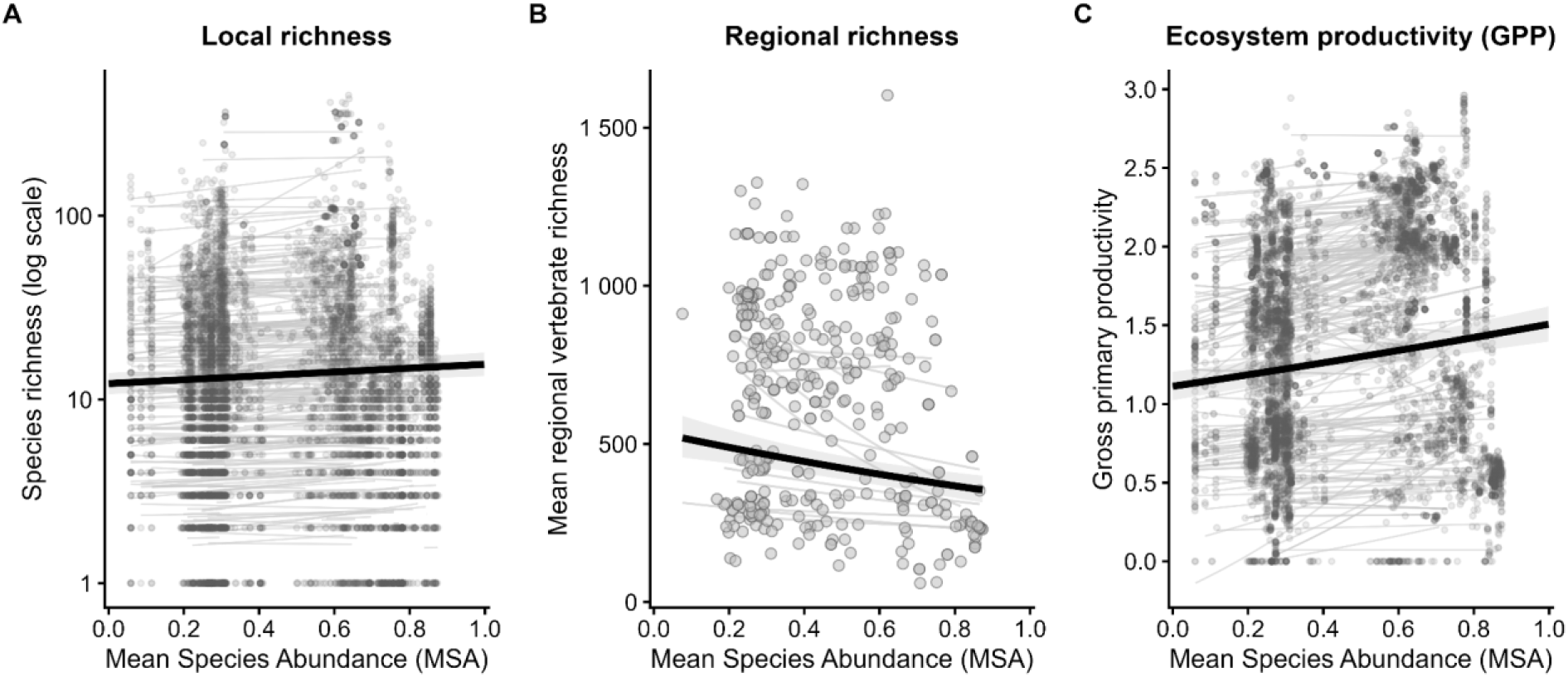
Overall relationships between Mean Species Abundance (MSA) and ecological outcomes. Relationships between MSA and (A) local species richness, (B) regional richness, and (C) ecosystem productivity (DHI-GPP). Points represent observations, black lines represent the overall fitted relationships, and light grey lines represent within-study fits for local richness and ecosystem productivity and within-biome fits for regional richness. Shaded areas around the overall fits represent 95% confidence intervals.

**Figure 3.**
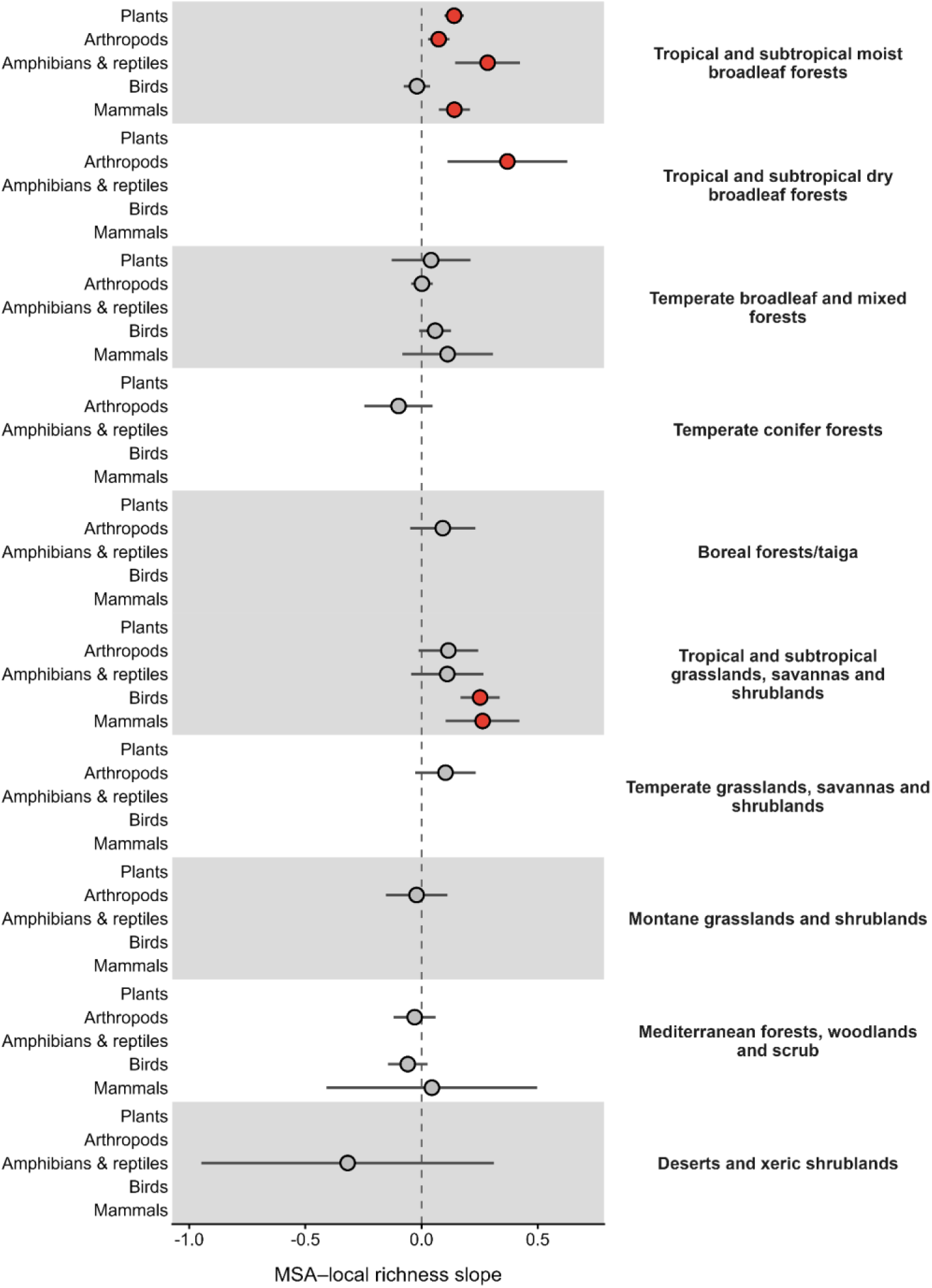
Variation in the relationship between MSA and local species richness across biomes and taxonomic groups. Points represent estimated MSA–richness slopes for each biome–taxon combination, and horizontal lines represent 95% confidence intervals. Red points indicate significant positive relationships, blue points indicate significant negative relationships, and grey points indicate non-significant relationships. Only biome–taxon with more than 10 sampling locations were included in the analyses.

The association between MSA and local species richness remained positive only for plants at high NDVI values, whereas it reversed for birds at low values of NDVI (figure 4, table S3). EVI altered the association between MSA and local richness only for plants (figure 4). We also found that landscape structure altered the magnitude and direction of this association. While increasing forest cover increased the positive association between MSA and local bird richness, it decreased the association between MSA and local mammal richness. Forest cover did not significantly change the MSA–richness association for other taxonomic groups (figure 4). Additionally, increasing habitat proximity also decreased the association between MSA and local mammal richness, yet it did not significantly alter the association for the other taxonomic groups.

**Figure 4.**
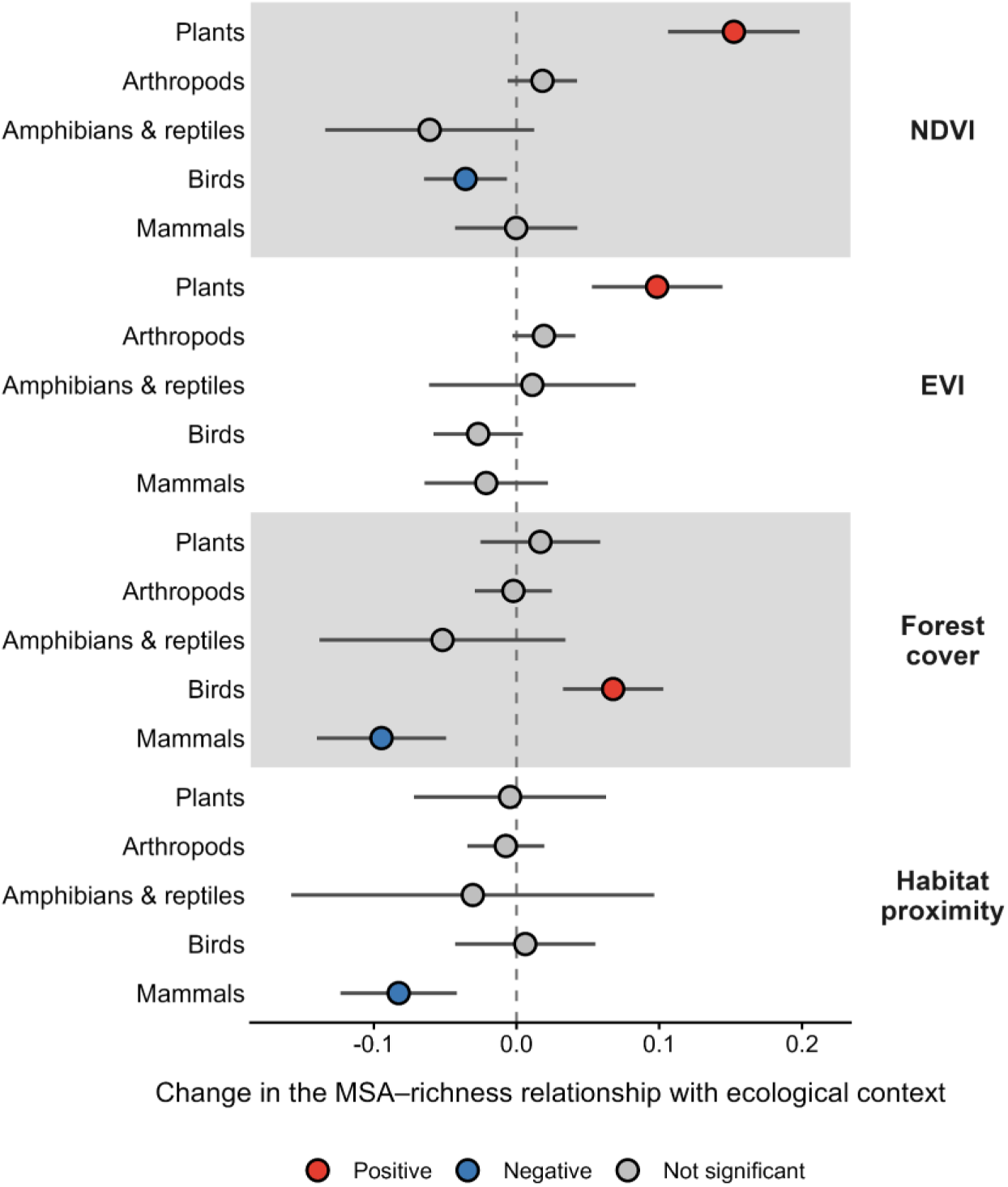
Effects of ecological context on the relationship between MSA and local species richness across taxonomic groups. Points show how the MSA–richness relationship changes with increasing NDVI, EVI, forest cover, or habitat proximity, and horizontal lines represent 95% confidence intervals. Positive estimates indicate that the MSA–richness relationship becomes more positive as the contextual variable increases, whereas negative estimates indicate that it becomes more negative. Red and blue points indicate significant positive and negative changes, respectively, and grey points indicate non-significant changes. Ecological-context variables were standardized before analysis.

The association with regional richness remained negative (figure 2B) only for temperate conifer and boreal forests (figure 5, table S3), but this association became positive for tropical and subtropical dry forests, tropical and subtropical grasslands, savannas, and shrublands, temperate broadleaf and mixed forests, and montane grasslands (figure 5). For the remaining biomes, we found no significant association between MSA and regional richness. Vegetation conditions also altered the direction of this association, with increasing NDVI and EVI favoring a stronger positive association between MSA and regional richness (figure 6). Conversely, increasing habitat proximity, favored a negative association between MSA and regional richness (figure 6).

**Figure 5.**
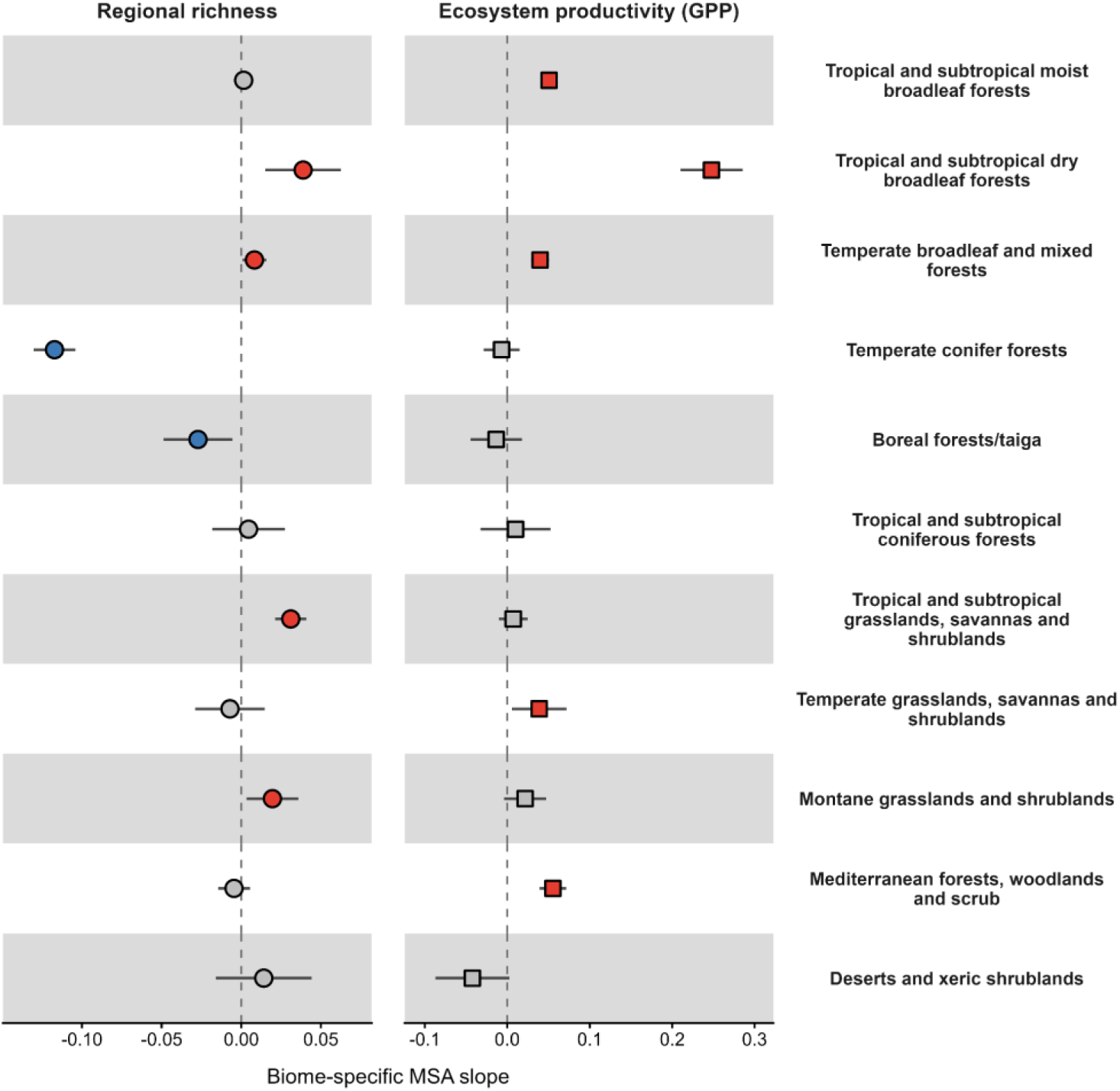
Variation in the relationships between MSA and regional species richness and ecosystem productivity across biomes. Points represent biome-specific MSA slopes for regional richness and ecosystem productivity (DHI-GPP), and horizontal lines represent 95% confidence intervals. Red symbols indicate significant positive relationships, blue symbols indicate significant negative relationships, and grey symbols indicate non-significant relationships.

**Figure 6.**
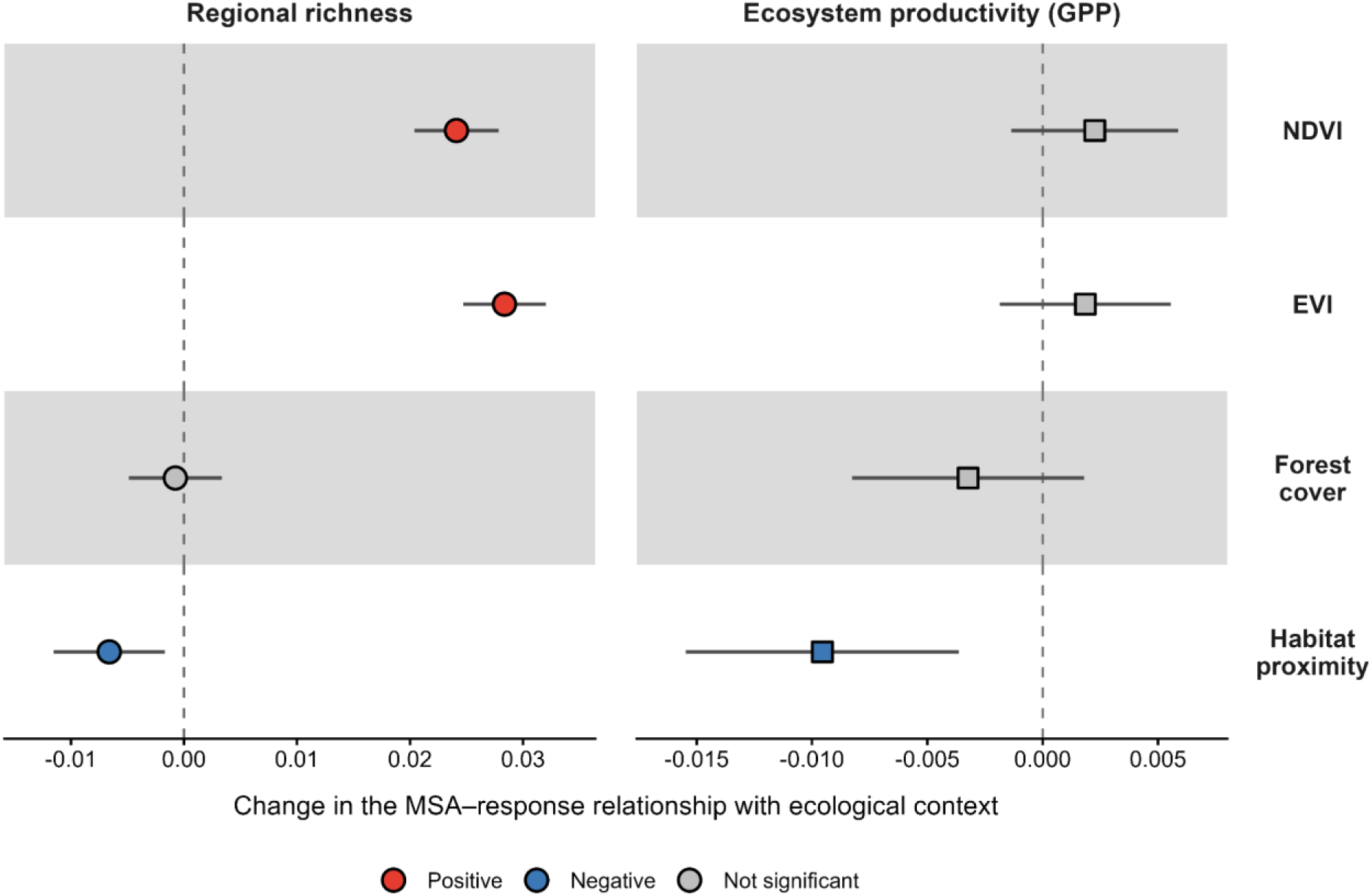
Effects of ecological context on the relationships between MSA and regional species richness and ecosystem productivity (DHI-GPP). Points show how the MSA–response relationship changes with increasing NDVI, EVI, forest cover, or habitat proximity, and horizontal lines represent 95% confidence intervals. Positive estimates indicate that the MSA– response relationship becomes more positive as the contextual variable increases, whereas negative estimates indicate that it becomes more negative. Red and blue symbols indicate significant positive and negative changes, respectively, and grey symbols indicate non-significant changes. Ecological-context variables were standardized before analysis.

MSA and ecosystem productivity generally increased in tandem (figure 1C). This relationship was retained in five of the 11 studied biomes: tropical and subtropical moist and dry broadleaf forests, temperate broadleaf and mixed forests, temperate grasslands, and Mediterranean forests. However, only habitat proximity altered the association between MSA and ecosystem productivity, as decreasing habitat proximity favored a negative association between MSA and ecosystem productivity (figure 6). The other ecological context variables did not significantly alter the association between MSA and ecosystem productivity.

At a local scale, we found that including within- and between-study abundance did not change most of the associations between MSA and local species richness (Δ estimates in table S3). The main changes occurred for models including landscape structure. In particular, the effects of forest cover on the MSA–richness relationship were substantially reduced after accounting for abundance, whereas associations involving vegetation conditions were comparatively robust (table S3). This indicates that some of the estimated landscape-structure effects on the MSA–richness relationship are sensitive to variation in abundance across sites.

## 4. Discussion

Although MSA captures some global variation in ecological outcomes, the magnitude and, in some cases, the direction of its associations with these outcomes vary across ecological contexts and taxonomic groups. The representativeness of MSA as a proxy for other dimensions of biodiversity and ecosystem properties depends on whether these relationships are transferable or conditional. MSA is transferable when its association with a given ecological outcome remains relatively consistent across ecological contexts. In contrast, MSA is context-dependent when ecological context substantially modifies this association. This conditionality emerges in two ways. First, ecological context can alter the direction of the relationship, making comparisons particularly challenging. For example, high MSA, indicating high biodiversity intactness, can occur in regions with comparatively low absolute species richness or productivity. Second, ecological context can alter the magnitude of the relationship, such that an overall association between MSA and an ecological outcome may weaken or disappear in specific contexts or taxonomic groups.

### 4.1. Relationships between MSA and ecological outcomes vary across ecological contexts and taxonomic groups

Simple models that ignored ecological context showed that biodiversity intactness increases alongside local species richness and ecosystem productivity. However, associations between MSA and ecosystems performance weakened, disappeared, or, in some cases, reversed direction when accounting for ecological context. Context dependence challenges the assumption that a common MSA response can be interpreted similarly across diverse environments (Raymond *et al* 2026). For example, MSA was positively associated with local species richness in several tropical and subtropical biomes, whereas this relationship was weak or absent in temperate biomes. Thus, similar changes in MSA do not necessarily correspond to similar changes in local species richness across ecological contexts.

These findings align with previous studies showing that habitat loss reduces local species richness more in protected areas than in degraded landscapes (Crouzeilles *et al* 2016, Betts *et al* 2017, Li *et al* 2024, Gonçalves-Souza *et al* 2025a) and more in tropical biomes than in higher-latitude temperate biomes (Newbold *et al* 2020b). As such, generalizing biodiversity responses across contrasting landscapes obscures important ecological differences. This issue is also relevant to the parameterization of MSA in GLOBIO. MSA values for a given land use are derived from pressure-impact relationships taken from multiple studies and are then combined with other spatially varying pressures, such as climate change and habitat fragmentation (Schipper *et al* 2020). Although these additional pressures introduce regional variation in MSA, the response assigned to a given land use does not explicitly vary among ecological contexts. Applying these values across contrasting regions potentially obscures differences in the magnitude or direction of ecological responses (Newbold *et al* 2015, VanderWilde and Newell 2021, Roeder and Utz 2025).

The influence of ecological context on the representativeness of MSA also differed among taxonomic groups, with context-dependent relationships for plants, birds, and mammals but little evidence of systematic moderation for arthropods and amphibians and reptiles. Collectively, these results reinforce the limitations of interpreting MSA as a universal proxy for species richness across taxonomic groups, an issue already recognized in the development and application of MSA-based models (Alkemade *et al* 2009, Schipper *et al* 2020) and in other biodiversity metrics (Heino 2010, Brown and Williams 2016). MSA was parameterized using datasets on plants, birds, and mammals, and the global projections that provide pixel-scale MSA values used different sets of pressure–impact drivers, with an unbalanced representation of these drivers across the taxonomic groups studied (Schipper *et al* 2020). Taxonomic groups can respond differently to environmental pressures, and the congruence among these responses can vary with taxonomic group and spatial scale (Piano *et al* 2020, Newbold *et al* 2020a). Moreover, the MSA–richness relationship may be transferable for some taxonomic groups but context-dependent for others.

Previous studies have used MSA as an indicator of cumulative human disturbance across space or time, interpreting it as an aggregate measure of biodiversity intactness and of how fragmentation, climate change, and nutrient pollution affect biodiversity across regions (Liao *et al* 2024, Schipper *et al* 2025, Ambrósio *et al* 2024). However, these broader-scale assessments are generally projected from relative, local-scale metrics. Our findings provide an empirical evaluation of when a relative metric such as MSA corresponds to absolute ecological outcomes across different spatial scales and response variables. In general, MSA showed a more consistent association with ecosystem productivity than with regional species richness, which is more context dependent. The contrasting spatial scales of regional species richness and MSA may also contribute to their observed relationship, because regional richness integrates biodiversity over a much broader area than the local-scale MSA estimates.

### 4.2. How ecological context can guide the interpretation and use of MSA in LCA

MSA is useful but it comes with caveats regarding ecological context, taxonomic group, and spatial scale. These dimensions can help identify how MSA should be interpreted and when its interpretation requires additional qualification in applications such as risk assessment, LCA, and certification (Vrasdonk *et al* 2019a, Myllyviita *et al* 2019, Bromwich *et al* 2026). Accordingly, we propose a decision tree with three branches that guide the use and interpretation of MSA by practitioners (figure 7).

**Figure 7.**
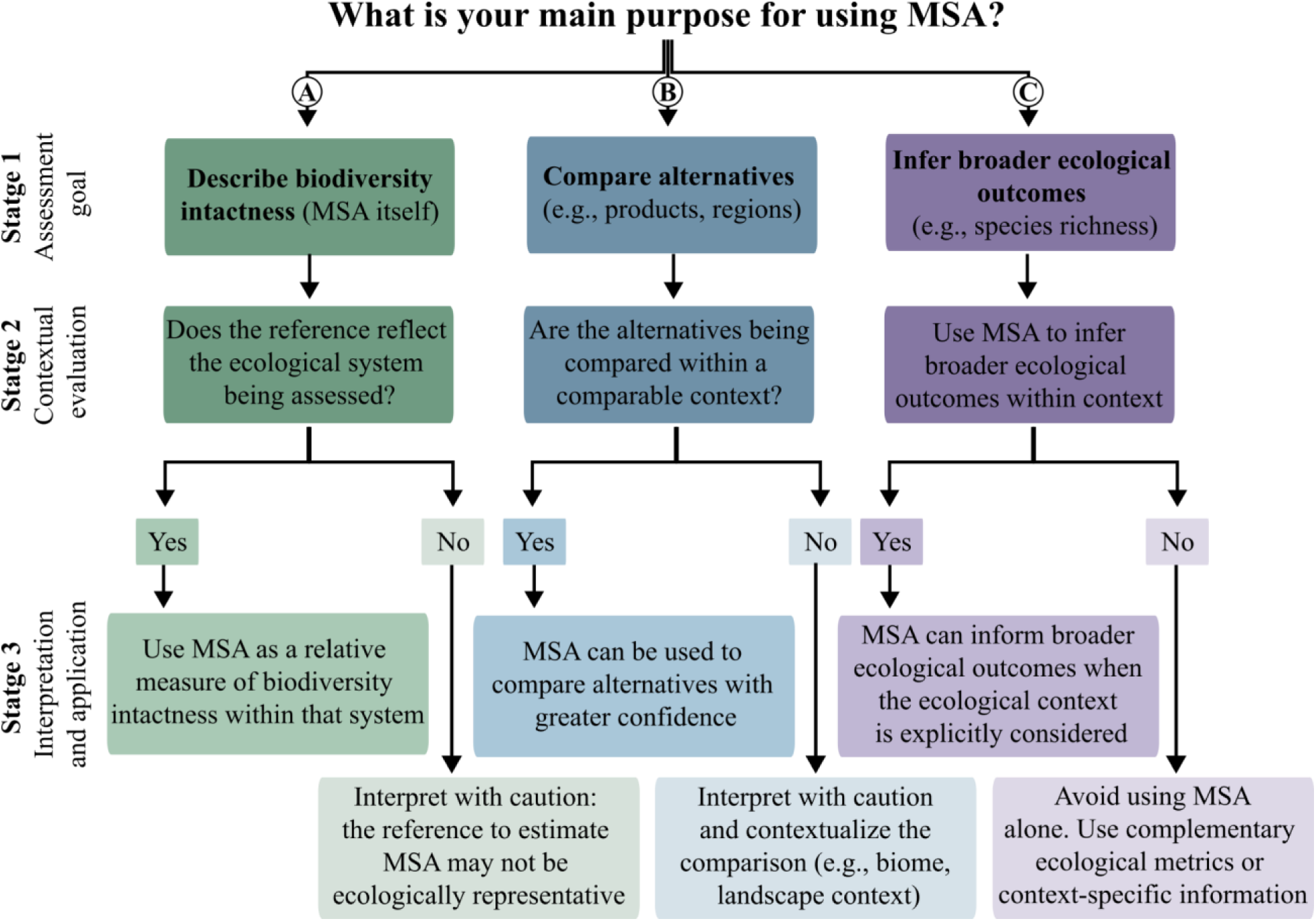
Decision guide for the interpretation and use of Mean Species Abundance (MSA). The decision tree separates three assessment goals: (A) describing biodiversity intactness, (B) comparing alternative systems, and (C) inferring broader ecological outcomes. For each goal, the decision tree evaluates whether the relevant reference condition or ecological context is appropriate for interpreting and applying MSA. The need for contextual information increases when moving from direct interpretation of biodiversity intactness toward comparisons among systems and inference of broader ecological outcomes.

The first step is to define the purpose for which MSA is being used: i) describing biodiversity intactness, ii) comparing alternative systems, such as products or regions, or iii) inferring broader ecological outcomes. Moving from the first to the second and third purposes requires progressively more information beyond the MSA value itself, because comparisons and ecological outcomes depend increasingly on context.

For assessments aimed at describing biodiversity intactness, the main consideration is whether the reference condition reflects the ecosystem being assessed (figure 7A). Since observations of human-free reference conditions are rare, studies often approximate them using relatively undisturbed or semi-natural habitats. Sometimes managed systems, such pastures are used as the reference (Vrasdonk *et al* 2019b). When the reference accurately represents the ecosystem being assessed, MSA can be interpreted as a relative measure of biodiversity intactness. Otherwise, greater caution is warranted when using MSA (figure 7A).

For comparisons among alternative systems, such as different production systems or the same product produced in different regions, the relevant question is whether those alternatives occur under comparable ecological contexts (figure 7B). Systems with similar landscape structures or management contexts provide a justifiable basis for interpreting differences in MSA. Comparisons across substantially different ecological contexts require greater caution (Alkemade *et al* 2009, Schipper *et al* 2020). In these cases, the ecological differences between systems should be explicitly reported so that the limitations of the comparison are transparent.

Ecological context needs to be considered when using MSA to infer ecological outcomes beyond biodiversity intactness (figure 7C). Regionalized biodiversity assessment has been proposed as an improvement over global-average characterization because biodiversity responses depend strongly on where impacts occur (de Baan *et al* 2013, Schipper *et al* 2025). Our results extend this reasoning by showing that ecological context can also modify how MSA relates to local species richness, regional species richness, and ecosystem productivity. Accordingly, interpretations that extend MSA beyond biodiversity intactness should consider relevant ecological-context, taxonomic, and spatial information. When this information is unavailable or comparisons span highly contrasting ecological systems, MSA should be considered alongside complementary ecological metrics to holistically capture ecosystem impacts (Kuipers *et al* 2025, Mendez Angarita *et al* 2025).

### 4.3. Conclusion and implications for future applications of MSA

Our findings demonstrate that the interpretation of MSA becomes conditional as it is extended from biodiversity intactness itself to comparisons among systems and to broader ecological scenarios. Ecological context, taxonomic group, and spatial scale influence whether the relationships between MSA and different ecological outcomes are transferable across systems, which ultimately influences whether MSA can be appropriately interpreted as an endpoint measure of biodiversity intactness. We provide a guide to help future studies identify when MSA can be interpreted on its own and when complementary metrics are needed. These findings reinforce the need for caution when treating MSA as an aggregate metric, particularly in LCA, global policy frameworks, corporate biodiversity accounting, and forecasting. By identifying the conditions under which MSA is transferable or context dependent, we define operational boundaries in supporting more sustainable production and consumption.

## Notes

### Competing Interest Statement

The authors have declared no competing interest.

## References

Alkemade R, van Oorschot M, Miles L, Nellemann C, Bakkenes M and ten Brink B 2009 GLOBIO3: A Framework to Investigate Options for Reducing Global Terrestrial Biodiversity Loss Ecosystems 12 374–90 Online: 10.1007/s10021-009-9229-5

Ambrósio G, Doelman J C, Schipper A M, Stehfest E and van Vuuren D 2024 Global sustainability scenarios lead to regionally different outcomes for terrestrial biodiversity Environ. Res. Lett. 19 104055 Online: 10.1088/1748-9326/ad73eb

Arets E J M M, Verwer C C and Alkemade J R M 2014 Meta-analysis of the effect of global warming on local species richness Report 34 Online: https://research.wur.nl/en/publications/meta-analysis-of-the-effect-of-global-warming-on-local-species-ri/

Avila-Ortega D I, Jørgensen P S, Cornell S E, Moran D D and Engstrom G 2025 Accounting for biodiversity impacts of consumption and production: current gaps and frontiers. EcoEvoRxiv Online: https://ecoevorxiv.org/repository/view/9107/

de Baan L, Mutel C L, Curran M, Hellweg S and Koellner T 2013 Land Use in Life Cycle Assessment: Global Characterization Factors Based on Regional and Global Potential Species Extinction Environ. Sci. Technol. 47 9281–90 Online: 10.1021/es400592q

Bao S, Wutzler T, Koirala S, Cuntz M, Ibrom A, Besnard S, Walther S, Šigut L, Moreno A, Weber U, Wohlfahrt G, Cleverly J, Migliavacca M, Woodgate W, Merbold L, Veenendaal E and Carvalhais N 2022 Environment-sensitivity functions for gross primary productivity in light use efficiency models Agric. For. Meteorol. 312 108708 Online: https://www.sciencedirect.com/science/article/pii/S0168192321003944

Beer C, Reichstein M, Tomelleri E, Ciais P, Jung M, Carvalhais N, Rödenbeck C, Arain M A, Baldocchi D, Bonan G B, Bondeau A, Cescatti A, Lasslop G, Lindroth A, Lomas M, Luyssaert S, Margolis H, Oleson K W, Roupsard O, Veenendaal E, Viovy N, Williams C, Woodward F I and Papale D 2010 Terrestrial Gross Carbon Dioxide Uptake: Global Distribution and Covariation with Climate Science 329 834–8 Online: https://www.science.org/doi/10.1126/science.1184984

Betts M G, Wolf C, Ripple W J, Phalan B, Millers K A, Duarte A, Butchart S H M and Levi T 2017 Global forest loss disproportionately erodes biodiversity in intact landscapes Nature 547 441–4 Online: https://www.nature.com/articles/nature23285

Bevan P A, Ferreira G B, Ingram D J, Rowcliffe M, Young L, Freeman R and Jones K E 2025 Regional Biomes outperform broader spatial units in capturing biodiversity responses to land-use change Ecography 2025 e07318 Online: https://onlinelibrary.wiley.com/doi/abs/10.1111/ecog.07318

Bromwich T, White T B, Bouchez A, Hawkins I, zu Ermgassen S, Bull J, Bartlett H, Bennun L, Biggs E, Booth H, Clark M, El Geneidy S, Prescott G W, Sonter L J, Starkey M and Milner-Gulland E J 2026 Navigating uncertainty in life cycle assessment-based approaches to biodiversity footprinting Methods Ecol. Evol. 17 1387–404 Online: https://onlinelibrary.wiley.com/doi/abs/10.1111/2041-210x.70001

Brooks M E, Kristensen K, Benthem K J, van, Magnusson A, Berg C W, Nielsen A, Skaug H J, Mächler M and Bolker B M 2017 glmmTMB Balances Speed and Flexibility Among Packages for Zero-inflated Generalized Linear Mixed Modeling R J. 9 378 Online: https://journal.r-project.org/archive/2017/RJ-2017-066/index.html

Brown E D and Williams B K 2016 Ecological integrity assessment as a metric of biodiversity: are we measuring what we say we are? Biodivers. Conserv. 25 1011–35 Online: 10.1007/s10531-016-1111-0

Burgess N D, Ali N, Bedford J, Bhola N, Brooks S, Cierna A, Correa R, Harris M, Hargey A, Hughes J, McDermott-Long O, Miles L, Ravilious C, Rodrigues A R, van Soesbergen A, Sihvonen H, Seager A, Swindell L, Vukelic M, Durán A P, Green J M H, West C, Weatherdon L V, Hawkins F, Brooks T M, Kingston N and Butchart S H M 2024 Global Metrics for Terrestrial Biodiversity Annu. Rev. Environ. Resour. 49 673–709 Online: https://www.annualreviews.org/content/journals/10.1146/annurev-environ-121522-045106

Cardinale B J, Duffy J E, Gonzalez A, Hooper D U, Perrings C, Venail P, Narwani A, Mace G M, Tilman D, Wardle D A, Kinzig A P, Daily G C, Loreau M, Grace J B, Larigauderie A, Srivastava D S and Naeem S 2012 Biodiversity loss and its impact on humanity Nature 486 59–67 Online: https://www.nature.com/articles/nature11148

Chase J M, Blowes S A, Knight T M, Gerstner K and May F 2020 Ecosystem decay exacerbates biodiversity loss with habitat loss Nature 584 238–43 Online: https://www.nature.com/articles/s41586-020-2531-2

Clements H S, Do Linh San E, Hempson G, Linden B, Maritz B, Monadjem A, Reynolds C, Siebert F, Stevens N, Biggs R, De Vos A, Blanchard R, Child M, Esler K J, Hamann M, Loft T, Reyers B, Selomane O, Skowno A L, Tshoke T, Abdoulaye D, Aebischer T, Aguirre-Gutiérrez J, Alexander G J, Ali A H, Allan D G, Amoako E E, Angedakin S, Aruna E, Avenant N L, Badjedjea G, Bakayoko A, Bamba-kaya A, Bates M F, Bates P J J, Belmain S R, Bennitt E, Bradley J, Brewster C A, Brown M B, Brown M, Bryja J, Butynski T M, Carvalho F, Channing A, Chapman C A, Cohen C, Cords M, Cramer J D, Cronk N, Cunneyworth P M K, Dalerum F, Danquah E, Davies-Mostert H T, de Blocq A D, De Jong Y A, Demos T C, Denys C, Djagoun C A M S, Doherty-Bone T M, Drouilly M, du Toit J T, Ehlers Smith D A, Ehlers Smith Y C, Eiseb S J, Fashing P J, Ferguson A W, Fernández-García J M, Finckh M, Fischer C, Gandiwa E, Gaubert P, Gaugris J Y, Gibbs D J, Gilchrist J S, Gil-Sánchez J M, Githitho A N, Goodman P S, Granjon L, Grobler J P, Gumbi B C, Gvozdik V, Harvey J, Hauptfleisch M, Hayder F, Hema E M, Herbst M, Houngbédji M, Huntley B J, Hutterer R, Ivande S T, Jackson K, Jongsma G F M, Juste J, Kadjo B, Kaleme P K, Kamugisha E, Kaplin B A, et al 2024 The bii4africa dataset of faunal and floral population intactness estimates across Africa’s major land uses Sci. Data 11 191 Online: https://www.nature.com/articles/s41597-023-02832-6

Crouzeilles R, Curran M, Ferreira M S, Lindenmayer D B, Grelle C E V and Rey Benayas J M 2016 A global meta-analysis on the ecological drivers of forest restoration success Nat. Commun. 7 11666 Online: https://www.nature.com/articles/ncomms11666

Damiani M, Sinkko T, Caldeira C, Tosches D, Robuchon M and Sala S 2023 Critical review of methods and models for biodiversity impact assessment and their applicability in the LCA context Environ. Impact Assess. Rev. 101 107134 Online: https://www.sciencedirect.com/science/article/pii/S0195925523001002

Dasgupta S, Blankespoor B and Wheeler D 2026 Toward better conservation: a spatial analysis of species occurrence data from the Global Biodiversity Information Facility Earth Syst. Sci. Data 18 2371–95 Online: https://essd.copernicus.org/articles/18/2371/2026/

Díaz S, Fargione J, Iii F S C and Tilman D 2006 Biodiversity Loss Threatens Human Well-Being PLOS Biol. 4 e277 Online: https://journals.plos.org/plosbiology/article?id=10.1371/journal.pbio.0040277

Dıaz S and Cabido M 2001 Vive la différence: plant functional diversity matters to ecosystem processes Trends Ecol. Evol. 16 646–55 Online: https://www.sciencedirect.com/science/article/pii/S0169534701022832

Dürr J, Dietz T and Börner J 2026 Telecoupling of governance systems under the EU deforestation regulation: Scenarios for the Brazilian beef and soy value chains For. Policy Econ. 185 103725 Online: https://www.sciencedirect.com/science/article/pii/S1389934126000304

El-Gabbas A 2026 A Global, Taxon-Stratified, High-Resolution Sampling-Effort Dataset From GBIF for Bias-Aware Ecological Modelling Divers. Distrib. 32 e70205 Online: https://onlinelibrary.wiley.com/doi/abs/10.1111/ddi.70205

Etard A and Newbold T 2024 Species-level correlates of land-use responses and climate-change sensitivity in terrestrial vertebrates Conserv. Biol. 38 e14208 Online: https://onlinelibrary.wiley.com/doi/abs/10.1111/cobi.14208

Feld C K, Martins da Silva P, Paulo Sousa J, De Bello F, Bugter R, Grandin U, Hering D, Lavorel S, Mountford O, Pardo I, Pärtel M, Römbke J, Sandin L, Bruce Jones K and Harrison P 2009 Indicators of biodiversity and ecosystem services: a synthesis across ecosystems and spatial scales Oikos 118 1862–71 Online: https://onlinelibrary.wiley.com/doi/abs/10.1111/j.1600-0706.2009.17860.x

Gonçalves-Souza T, Chase J M, Haddad N M, Vancine M H, Didham R K, Melo F L P, Aizen M A, Bernard E, Chiarello A G, Faria D, Gibb H, Lima M G, Magnago L F S, Mariano-Neto E, Nogueira A A, Nemésio A, Passamani M, Pinho B X, Rocha-Santos L, Rodrigues R C, Safar N V H, Santos B A, Soto-Werschitz A, Tabarelli M, Uehara-Prado M, Vasconcelos H L, Vieira S and Sanders N J 2025a Species turnover does not rescue biodiversity in fragmented landscapes Nature 640 702–6

Gonçalves-Souza T, Vancine M H, Sanders N J, Haddad N M, Cortinhas L, Aase A L T O, de Aguiar W M, Aizen M A, Arroyo-Rodríguez V, Baz A, Benchimol M, Bernard E, Bertotto T J, Bispo A A, Bogoni J A, Boldorini G X, Bragagnolo C, Brosi B, Cantalice A S, do Carmo R F R, Cazeta E, Chiarello A G, de la Sancha N U, Didham R K, Faria D, Filgueiras B, Figueira J E C, Galvão G A, Garey M V, Gibb H, Gómez-Martínez C, González E, de Gusmão R A F, Henry M, de Jesus S, Kloss T G, Lázaro A, Leandro-Silva V, de Lima M G, da Silva Lima I, Lins-e-Silva A C B, Nally R M, Magalhães A R, Magnago L F S, Manu S, Mariano-Neto E, Mbora D N M, Melo F P L, Mutua M N, Neckel-Oliveira S, Nemésio A, Nogueira A A, Oliveira P M D A, Pádua D G, Paes L, de Paiva A B, Passamani M, Pena J C, Peres C A, Pinho B X, Pons J-M, Prasniewski V M, Reiniö J, dos Santos Rocha M, Rocha-Santos L, Rodal M J, Rodrigues R C, Safar N V H, Salomão R P, Santos B A, Santos M N, dos Santos J P, Savilaakso S, Schaefer C E G R, Silva M A M, da Silva F R, Silva R J, Simonelli M, Soto-Werschitz A, Stireman III J O, Storck-Tonon D, Szinwelski N, Tabarelli M, Teixeira C P, Totland Ø, Uehara-Prado M, Vaz-de-Mello F Z, Vasconcelos H L, Vieira S A and Chase J M 2025b LandFrag: A Dataset to Investigate the Effects of Forest Loss and Fragmentation on Biodiversity Glob. Ecol. Biogeogr. 34 e70015 Online: https://onlinelibrary.wiley.com/doi/abs/10.1111/geb.70015

Gounaridis D, Newell J P and Goodspeed R 2020 The impact of urban sprawl on forest landscapes in Southeast Michigan, 1985–2015 Landsc. Ecol. 35 1975–93 Online: https://link.springer.com/10.1007/s10980-020-01075-9

Hardin J and Hilbe J 2018 Generalized Linear Models and Extensions (Texas: Stata Press)

Hayashi K 2020 Inconsistencies between regional- and field-scale biodiversity indicators within life cycle assessment: the case of rice production systems in Japan Int. J. Life Cycle Assess. 25 1278–89 Online: 10.1007/s11367-020-01749-1

Heino J 2010 Are indicator groups and cross-taxon congruence useful for predicting biodiversity in aquatic ecosystems? Ecol. Indic. 10 112–7 Online: https://www.sciencedirect.com/science/article/pii/S1470160X09000855

Hellweg S, Benetto E, Huijbregts M A J, Verones F and Wood R 2023 Life-cycle assessment to guide solutions for the triple planetary crisis Nat. Rev. Earth Environ. 4 471–86 Online: https://www.nature.com/articles/s43017-023-00449-2

Hillebrand H, Blasius B, Borer E T, Chase J M, Downing J A, Eriksson B K, Filstrup C T, Harpole W S, Hodapp D, Larsen S, Lewandowska A M, Seabloom E W, Van de Waal D B and Ryabov A B 2018 Biodiversity change is uncoupled from species richness trends: Consequences for conservation and monitoring J. Appl. Ecol. 55 169–84 Online: https://onlinelibrary.wiley.com/doi/abs/10.1111/1365-2664.12959

Hudson L N, Newbold T, Contu S, Hill S L L, Lysenko I, De Palma A, Phillips H R P, Senior R A, Bennett D J, Booth H, Choimes A, Correia D L P, Day J, Echeverría-Londoño S, Garon M, Harrison M L K, Ingram D J, Jung M, Kemp V, Kirkpatrick L, Martin C D, Pan Y, White H J, Aben J, Abrahamczyk S, Adum G B, Aguilar-Barquero V, Aizen M A, Ancrenaz M, Arbeláez-Cortés E, Armbrecht I, Azhar B, Azpiroz A B, Baeten L, Báldi A, Banks J E, Barlow J, Batáry P, Bates A J, Bayne E M, Beja P, Berg Å, Berry N J, Bicknell J E, Bihn J H, Böhning-Gaese K, Boekhout T, Boutin C, Bouyer J, Brearley F Q, Brito I, Brunet J, Buczkowski G, Buscardo E, Cabra-García J, Calviño-Cancela M, Cameron S A, Cancello E M, Carrijo T F, Carvalho A L, Castro H, Castro-Luna A A, Cerda R, Cerezo A, Chauvat M, Clarke F M, Cleary D F R, Connop S P, D’Aniello B, da Silva P G, Darvill B, Dauber J, Dejean A, Diekötter T, Dominguez-Haydar Y, Dormann C F, Dumont B, Dures S G, Dynesius M, Edenius L, Elek Z, Entling M H, Farwig N, Fayle T M, Felicioli A, Felton A M, Ficetola G F, Filgueiras B K C, Fonte S J, Fraser L H, Fukuda D, Furlani D, Ganzhorn J U, Garden J G, Gheler-Costa C, Giordani P, Giordano S, Gottschalk M S, et al 2014 The PREDICTS database: a global database of how local terrestrial biodiversity responds to human impacts Ecol. Evol. 4 4701–35 Online: https://onlinelibrary.wiley.com/doi/abs/10.1002/ece3.1303

IPBES 2019 Global assessment report on biodiversity and ecosystem services of the Intergovernmental Science-Policy Platform on Biodiversity and Ecosystem Services (Zenodo) Online: https://zenodo.org/records/6417333

IUCN 2026 The IUCN Red List of Threatened Species Online: https://www.iucnredlist.org. Accessed in 2026-01-09.

Japan Aerospace Exploration Agency (FNF4) 2016 Global 25 m Resolution Forest/Non-Forest Map (FNF) Online: https://eolp.jaxa.jp/research/PALSAR_PALSAR-2_FNF_map.html

Jaureguiberry P, Titeux N, Wiemers M, Bowler D E, Coscieme L, Golden A S, Guerra C A, Jacob U, Takahashi Y, Settele J, Díaz S, Molnár Z and Purvis A 2022 The direct drivers of recent global anthropogenic biodiversity loss Sci. Adv. 8 eabm9982 Online: https://www.science.org/doi/full/10.1126/sciadv.abm9982

Jegen M 2024 Life cycle assessment: from industry to policy to politics Int. J. Life Cycle Assess. 29 597–606 Online: 10.1007/s11367-023-02273-8

Kissling W D, Ahumada J A, Bowser A, Fernandez M, Fernández N, García E A, Guralnick R P, Isaac N J B, Kelling S, Los W, McRae L, Mihoub J-B, Obst M, Santamaria M, Skidmore A K, Williams K J, Agosti D, Amariles D, Arvanitidis C, Bastin L, De Leo F, Egloff W, Elith J, Hobern D, Martin D, Pereira H M, Pesole G, Peterseil J, Saarenmaa H, Schigel D, Schmeller D S, Segata N, Turak E, Uhlir P F, Wee B and Hardisty A R 2018 Building essential biodiversity variables (EBVs) of species distribution and abundance at a global scale Biol. Rev. 93 600–25 Online: https://onlinelibrary.wiley.com/doi/abs/10.1111/brv.12359

Kuipers K JJ, Melki A, Morel S and Schipper A M 2025 Relationships between mean species abundance (MSA) and potentially disappeared fraction of species (PDF) are consistent but also uncertain Environ. Sustain. Indic. 26 100652 Online: https://www.sciencedirect.com/science/article/pii/S266597272500073X

Li G, Fang C, Watson J E M, Sun S, Qi W, Wang Z and Liu J 2024 Mixed effectiveness of global protected areas in resisting habitat loss Nat. Commun. 15 8389 Online: https://www.nature.com/articles/s41467-024-52693-9

Liao M, Zhang Z, Yan R and Bai K 2024 The Assessment of Biodiversity Changes and Sustainable Agricultural Development in The Beijing-Tianjin-Hebei Region of China Sustainability 16 5678 Online: https://www.mdpi.com/2071-1050/16/13/5678

Liu D, Silveira E M O, Razenkova E, Anand A, Dubinin M, Hobi M, Pidgeon A M and Radeloff V C 2026 Global dynamic habitat indices (DHIs) based on MODIS and VIIRS vegetation products Remote Sens. Environ. 332 115099 Online: https://www.sciencedirect.com/science/article/pii/S0034425725005036

Lüscher G, Nemecek T, Arndorfer M, Balázs K, Dennis P, Fjellstad W, Friedel J K, Gaillard G, Herzog F, Sarthou J-P, Stoyanova S, Wolfrum S and Jeanneret P 2017 Biodiversity assessment in LCA: a validation at field and farm scale in eight European regions Int. J. Life Cycle Assess. 22 1483–92 Online: 10.1007/s11367-017-1278-y

Marques A, Robuchon M, Hellweg S, Newbold T, Beher J, Bekker S, Essl F, Ehrlich D, Hill S, Jung M, Marquardt S, Rosa F, Rugani B, Suárez-Castro A F, Silva A P, Williams D R, Dubois G and Sala S 2021 A research perspective towards a more complete biodiversity footprint: a report from the World Biodiversity Forum Int. J. Life Cycle Assess. 26 238–43 Online: 10.1007/s11367-020-01846-1

McGarigal K, Cushman S and Ene E 2023 FRAGSTATS: Spatial Pattern Analysis Program for Categorical Maps.

McNellie M J, Oliver I, Dorrough J, Ferrier S, Newell G and Gibbons P 2020 Reference state and benchmark concepts for better biodiversity conservation in contemporary ecosystems Glob. Change Biol. 26 6702–14 Online: https://onlinelibrary.wiley.com/doi/abs/10.1111/gcb.15383

Mendez Angarita V Y, Larsen P B, Marcolin L and Di Marco M 2025 Reconciling Different Forms of Ecological Integrity to Aid the Kunming-Montreal Global Biodiversity Framework Conserv. Lett. 18 e13088 Online: https://onlinelibrary.wiley.com/doi/abs/10.1111/conl.13088

Myllyviita T, Sironen S, Saikku L, Holma A, Leskinen P and Palme U 2019 Assessing biodiversity impacts in life cycle assessment framework - Comparing approaches based on species richness and ecosystem indicators in the case of Finnish boreal forests J. Clean. Prod. 236 117641 Online: https://www.sciencedirect.com/science/article/pii/S0959652619324916

Naeem S, Chazdon R, Duffy J E, Prager C and Worm B 2016 Biodiversity and human well-being: an essential link for sustainable development Proc. R. Soc. B Biol. Sci. 283 20162091 Online: 10.1098/rspb.2016.2091

Neel M C, McGarigal K and Cushman S A 2004 Behavior of class-level landscape metrics across gradients of class aggregation and area Landsc. Ecol. 19 435–55 Online: 10.1023/B:LAND.0000030521.19856.cb

Newbold T, Bentley L F, Hill S L L, Edgar M J, Horton M, Su G, Şekercioğlu Ç H, Collen B and Purvis A 2020a Global effects of land use on biodiversity differ among functional groups Funct. Ecol. 34 684–93 Online: https://onlinelibrary.wiley.com/doi/abs/10.1111/1365-2435.13500

Newbold T, Hudson L N, Hill S L L, Contu S, Lysenko I, Senior R A, Börger L, Bennett D J, Choimes A, Collen B, Day J, De Palma A, Díaz S, Echeverria-Londoño S, Edgar M J, Feldman A, Garon M, Harrison M L K, Alhusseini T, Ingram D J, Itescu Y, Kattge J, Kemp V, Kirkpatrick L, Kleyer M, Correia D L P, Martin C D, Meiri S, Novosolov M, Pan Y, Phillips H R P, Purves D W, Robinson A, Simpson J, Tuck S L, Weiher E, White H J, Ewers R M, Mace G M, Scharlemann J P W and Purvis A 2015 Global effects of land use on local terrestrial biodiversity Nature 520 45–50 Online: https://www.nature.com/articles/nature14324

Newbold T, Oppenheimer P, Etard A and Williams J J 2020b Tropical and Mediterranean biodiversity is disproportionately sensitive to land-use and climate change Nat. Ecol. Evol. 4 1630–8 Online: https://www.nature.com/articles/s41559-020-01303-0

Pereira H M, Martins I S, Rosa I M D, Kim H, Leadley P, Popp A, van Vuuren D P, Hurtt G, Quoss L, Arneth A, Baisero D, Bakkenes M, Chaplin-Kramer R, Chini L, Di Marco M, Ferrier S, Fujimori S, Guerra C A, Harfoot M, Harwood T D, Hasegawa T, Haverd V, Havlík P, Hellweg S, Hilbers J P, Hill S L L, Hirata A, Hoskins A J, Humpenöder F, Janse J H, Jetz W, Johnson J A, Krause A, Leclère D, Matsui T, Meijer J R, Merow C, Obersteiner M, Ohashi H, De Palma A, Poulter B, Purvis A, Quesada B, Rondinini C, Schipper A M, Settele J, Sharp R, Stehfest E, Strassburg B B N, Takahashi K, Talluto L, Thuiller W, Titeux N, Visconti P, Ware C, Wolf F and Alkemade R 2024 Global trends and scenarios for terrestrial biodiversity and ecosystem services from 1900 to 2050 Science 384 458–65 Online: https://www.science.org/doi/10.1126/science.adn3441

Piano E, Souffreau C, Merckx T, Baardsen L F, Backeljau T, Bonte D, Brans K I, Cours M, Dahirel M, Debortoli N, Decaestecker E, Wolf K D, Engelen J M T, Fontaneto D, Gianuca A T, Govaert L, Hanashiro F T T, Higuti J, Lens L, Martens K, Matheve H, Matthysen E, Pinseel E, Sablon R, Schön I, Stoks R, Doninck K V, Dyck H V, Vanormelingen P, Wichelen J V, Vyverman W, Meester L D and Hendrickx F 2020 Urbanization drives cross-taxon declines in abundance and diversity at multiple spatial scales Glob. Change Biol. 26 1196–211 Online: https://onlinelibrary.wiley.com/doi/10.1111/gcb.14934

Radeloff V C, Dubinin M, Coops N C, Allen A M, Brooks T M, Clayton M K, Costa G C, Graham C H, Helmers D P, Ives A R, Kolesov D, Pidgeon A M, Rapacciuolo G, Razenkova E, Suttidate N, Young B E, Zhu L and Hobi M L 2019 The Dynamic Habitat Indices (DHIs) from MODIS and global biodiversity Remote Sens. Environ. 222 204–14 Online: https://www.sciencedirect.com/science/article/pii/S0034425718305625

Raymond J, van’t Veen H, Brown C, Byari M, Repo A, Alkemade R, Zeng Y and Rounsevell M 2026 Shifts in forest management intensity mediate biodiversity outcomes under contrasting socio-economic and climate scenarios Environ. Res. Lett. 21 174026 Online: 10.1088/1748-9326/ae9781

Regos A, Gonçalves J, Arenas-Castro S, Alcaraz-Segura D, Guisan A and Honrado J P 2022 Mainstreaming remotely sensed ecosystem functioning in ecological niche models Remote Sens. Ecol. Conserv. 8 431–47 Online: https://onlinelibrary.wiley.com/doi/abs/10.1002/rse2.255

Reich P B, Tilman D, Isbell F, Mueller K, Hobbie S E, Flynn D F B and Eisenhauer N 2012 Impacts of Biodiversity Loss Escalate Through Time as Redundancy Fades Science 336 589–92 Online: https://www.science.org/doi/10.1126/science.1217909

Roeder R and Utz S 2025 From Diversity to Confusion? The Challenge of Biodiversity Footprint Quantification Bus. Strategy Environ. 34 5887–900 Online: https://onlinelibrary.wiley.com/doi/10.1002/bse.4215

Sala S, Crenna E, Secchi M and Sanyé-Mengual E 2020 Environmental sustainability of European production and consumption assessed against planetary boundaries J. Environ. Manage. 269 110686 Online: https://www.sciencedirect.com/science/article/pii/S0301479720306186

Santini L, Belmaker J, Costello M J, Pereira H M, Rossberg A G, Schipper A M, Ceaușu S, Dornelas M, Hilbers J P, Hortal J, Huijbregts M A J, Navarro L M, Schiffers K H, Visconti P and Rondinini C 2017 Assessing the suitability of diversity metrics to detect biodiversity change Biol. Conserv. 213 341–50 Online: https://www.sciencedirect.com/science/article/pii/S0006320716303305

Schipper A M, Hilbers J P, Meijer J R, Antão L H, Benítez-López A, de Jonge M M J, Leemans L H, Scheper E, Alkemade R, Doelman J C, Mylius S, Stehfest E, van Vuuren D P, van Zeist W-J and Huijbregts M A J 2020 Projecting terrestrial biodiversity intactness with GLOBIO 4 Glob. Change Biol. 26 760–71 Online: https://onlinelibrary.wiley.com/doi/abs/10.1111/gcb.14848

Schipper A M, van der Marel M, Bakkenes M, Giesen P, van Oorschot M M P, Wilting H C, Huijbregts M A J and Marques A 2025 Impact factors for quantifying country-level terrestrial biodiversity intactness footprints (IBIF) Sci. Data 12 1660 Online: https://www.nature.com/articles/s41597-025-05946-1

Sidemo-Holm W, Alkan Olsson J and Smith H G 2026 Biodiversity at risk when businesses rely on narrow indicators J. Clean. Prod. 574 149032 Online: https://www.sciencedirect.com/science/article/pii/S0959652626015738

Song J, Chaudhary A, Tang R, Bai X and Yang Y 2025 Life-cycle biodiversity assessment of global crop production: Advances, framework, and outlooks J. Environ. Manage. 393 126934 Online: https://www.sciencedirect.com/science/article/pii/S030147972502910X

Stevenson S L, Watermeyer K, Caggiano G, Fulton E A, Ferrier S and Nicholson E 2021 Matching biodiversity indicators to policy needs Conserv. Biol. 35 522–32 Online: https://onlinelibrary.wiley.com/doi/abs/10.1111/cobi.13575

Teixeira R F M, Maia de Souza D, Curran M P, Antón A, Michelsen O and Milài Canals L 2016 Towards consensus on land use impacts on biodiversity in LCA: UNEP/SETAC Life Cycle Initiative preliminary recommendations based on expert contributions J. Clean. Prod. 112 4283–7 Online: https://www.sciencedirect.com/science/article/pii/S0959652615010495

VanderWilde C P and Newell J P 2021 Ecosystem services and life cycle assessment: A bibliometric review Resour. Conserv. Recycl. 169 105461 Online: https://www.sciencedirect.com/science/article/pii/S0921344921000689

Vrasdonk E, Palme U and Lennartsson T 2019a Reference situations for biodiversity in life cycle assessments: conceptual bridging between LCA and conservation biology Int. J. Life Cycle Assess. 24 1631–42 Online: 10.1007/s11367-019-01594-x

Vrasdonk E, Palme U and Lennartsson T 2019b Reference situations for biodiversity in life cycle assessments: conceptual bridging between LCA and conservation biology Int. J. Life Cycle Assess. 24 1631–42 Online: 10.1007/s11367-019-01594-x

Weinzettel J, Vačkářů D and Medková H 2019 Potential net primary production footprint of agriculture: A global trade analysis J. Ind. Ecol. 23 1133–42 Online: https://onlinelibrary.wiley.com/doi/abs/10.1111/jiec.12850

Winfree R, W. Fox J, Williams N M, Reilly J R and Cariveau D P 2015 Abundance of common species, not species richness, drives delivery of a real-world ecosystem service Ecol. Lett. 18 626–35 Online: https://onlinelibrary.wiley.com/doi/abs/10.1111/ele.12424

Wüest R O, Zimmermann N E, Zurell D, Alexander J M, Fritz S A, Hof C, Kreft H, Normand S, Cabral J S, Szekely E, Thuiller W, Wikelski M and Karger D N 2020 Macroecology in the age of Big Data – Where to go from here? J. Biogeogr. 47 1–12 Online: https://onlinelibrary.wiley.com/doi/abs/10.1111/jbi.13633

Zhang P, Seabloom E W, Foo J, MacDougall A S, Harpole W S, Adler P B, Hautier Y, Eisenhauer N, Spohn M, Bakker J D, Lekberg Y, Young A L, Carbutt C, Risch A C, Peri P L, Smith N G, Stevens C J, Prober S M, Knops J M H, Wardle G M, Dickman C R, Ebeling A, Roscher C, Martinson H M, Martina J P, Power S A, Niu Y, Ren Z, Du G, Virtanen R, Tognetti P, Tedder M J, Jentsch A, Catford J A and Borer E T 2025 Dominant species predict plant richness and biomass in global grasslands Nat. Ecol. Evol. 9 924–36 Online: https://www.nature.com/articles/s41559-025-02701-y

Zhang Y, Rasul K, Dorber M, Stadler K, Hertwich E G and Verones F 2026 Terrestrial biodiversity impacts of replacing beef with alternative-protein burger patties across ten European Union Countries J. Ind. Ecol. 30 1761–77 Online: 10.1007/s44498-026-00120-1

