## Supplementary Materials for "Ecological context shapes the interpretation of biodiversity intactness metrics"

**1. Supplementary methods**
**1.1. Pressure-response relationships to calculate MSA**
**1.2. Database and biome level information**
**1.3. Vegetation condition and ecosystem variables**
**1.4. Model equations**
**1.5. Sensitivity analysis**

**2. Figures and Tables**
**Figure S1**. Global distribution of the sampling locations used to estimate local species richness for each taxonomic group. Panels show plants, arthropods, amphibians & reptiles, birds, and mammals. Each point represents the geographic centroid of a sampled continuous forest or forest fragment included in the analysis. Point color indicates the source database: LandFrag (orange) and PREDICTS (purple).
**Table S1**. Description of biodiversity metrics, ecosystem variables, and ecological-context predictors used in this study. Variables are grouped by analytical role, ecological dimension, scale, spatial resolution, and data source (see also figure 1, main text).
**Table S2**. Overall relationships between Mean Species Abundance (MSA) and local and regional species richness, and ecosystem productivity (GPP). Estimates represent standardized MSA effects. For local richness and GPP, coefficients represent within-study MSA effects, whereas the regional richness coefficient represents the within-biome MSA effect after accounting for broad differences among biomes and repeated studies. Within-study and within-biome fits are represented by the light grey lines in Fig. 2 (main text).

**Table S3.** Sensitivity analysis of MSA–local species richness relationships using abundance adjusted models. Taxon-specific changes in the effect of ecological context on the MSA–local species richness relationship are shown for the main models and models additionally accounting for within- and between-study variation in total abundance. Estimates represent the change in the MSA slope associated with a 1-SD increase in ecological context. Δ estimate is the abundance-adjusted estimate minus the main-model estimate.

### 1. Supplementary methods:

#### 1.1. Pressure-response relationships to calculate MSA

Each raster cell contains a modelled MSA estimate derived from pressure–response relationships for land use, climate change, habitat fragmentation, atmospheric nitrogen deposition, road disturbance, and hunting. For each grid cell, GLOBIO combines the effects of different pressures by multiplying their MSA values; this calculation is performed separately for plants, birds, and mammals. The combined MSA in a raster cell is the arithmetic mean of the plant and vertebrate MSA estimates.

For climate, Schipper et al (2020) used the Fraction of Remaining Species (Nunez *et al* 2019) as a proxy for MSA.

#### 1.2. Database and biome level information

From the 301studies collected from the LandFrag (Gonçalves-Souza *et al* 2025b) and PREDICTS (Hudson *et al* 2014) databases, we compiled data for 4,279,945 individuals from 21,761 species across five taxonomic groups in 11 biomes. Collectively, the studies were conducted over an approximately 40-year period, although the year of sampling varied among studies. These included 10,120 arthropods (n = 140 studies), 7,016 plants (n = 42 studies), 3,277 birds (n = 56 studies), 689 amphibians and reptiles (n = 32 studies), and 659 mammals (n = 48 studies) species. The most studied biome was tropical and subtropical moist broadleaf forests (166 studies and 3,266 sampling locations), and the least studied were mangroves (8 studies and 14 sampling locations) and tundra (1 study and 10 sampling locations).

#### 1.3. Vegetation condition and ecosystem variables

NDVI is widely used to characterize vegetation greenness and cover (Xu *et al* 2023), whereas EVI is designed to retain greater sensitivity in densely vegetated areas and reduce both atmospheric and background-soil influences (Huete *et al* 2002). The cumulative DHI-GPP represents the sum of the median GPP values for each of the 46 eight-day time steps of the composite annual phenological cycle derived from 2013 to 2022. It therefore represents the typical annual cumulative productivity regime at each sampling location (Radeloff *et al* 2019, Liu *et al* 2026)

### 1.4. Model equations

For the negative-binomial species-richness models, the conditional mean was modeled as:

$$log(\mu_{ij}) = \beta_0 + \beta_1 X_{ij} + \beta_2 Z_{ij} + \beta_3 X_{ij} Z_{ij} + b_j$$

where $i$ indexes observations and $j$ indexes studies, $\mu_{ij} = E(Y_{ij}| b_j)$ is the expected species richness for observation $i$ in study $j$, $X_{ij}$ is the standardized MSA, $Z_{ij}$ is the ecological-context variable, $\beta_0$ is the overall intercept, $\beta_1$ and $\beta_2$ are the main effects of MSA and ecological-context, respectively, $\beta_3$ is the MSA × ecological-context interaction effect, and $b_j$ is the study-specific random intercept, assumed to follow $b_j = N(0, \sigma^2_{study})$.

For the Gaussian ecosystem productivity models, we modeled log-transformed DHI-GPP as:

$$log(1 + GPP_{ij}) = \beta_0 + \beta_1 X_{ij} + \beta_2 Z_{ij} + \beta_3 X_{ij} Z_{ij} + b_j + \varepsilon_{ij}$$

where the model terms are defined as above and the residual error was assumed to follow $\varepsilon_{ij} = N(0, \varepsilon^2)$.

### 1.5. Sensitivity analysis

Sampling intensity determines our ability to estimate the number of species sampled in a given fragment (Gotelli and Colwell 2001, Gonçalves-Souza *et al* 2025a). As such, we used the total number of individuals as the realized sample size describing sampling intensity. We included abundance using two different terms. First, the within-study abundance term allowed us to account for the influence of the number of individuals sampled on the number of species observed among fragments within the same study. Second, the between-study abundance term controlled for systematic differences among studies in aspects such as sampling method, taxonomic group, and sampling duration (see, e.g., Chase *et al* 2020). Similarly, previous studies have partitioned abundance into within- and between-study components in mixed models (van de Pol and Wright 2009) to examine how variation in abundance influences the estimated effects of other relevant fixed variables (Denomme-Brown *et al* 2021, Tomašových *et al* 2024, Byrnes and

Dee 2025). Here, we assessed how the inclusion of within- and between-study abundance affected the main effects of MSA and ecological context.

### 2. Figures and Tables

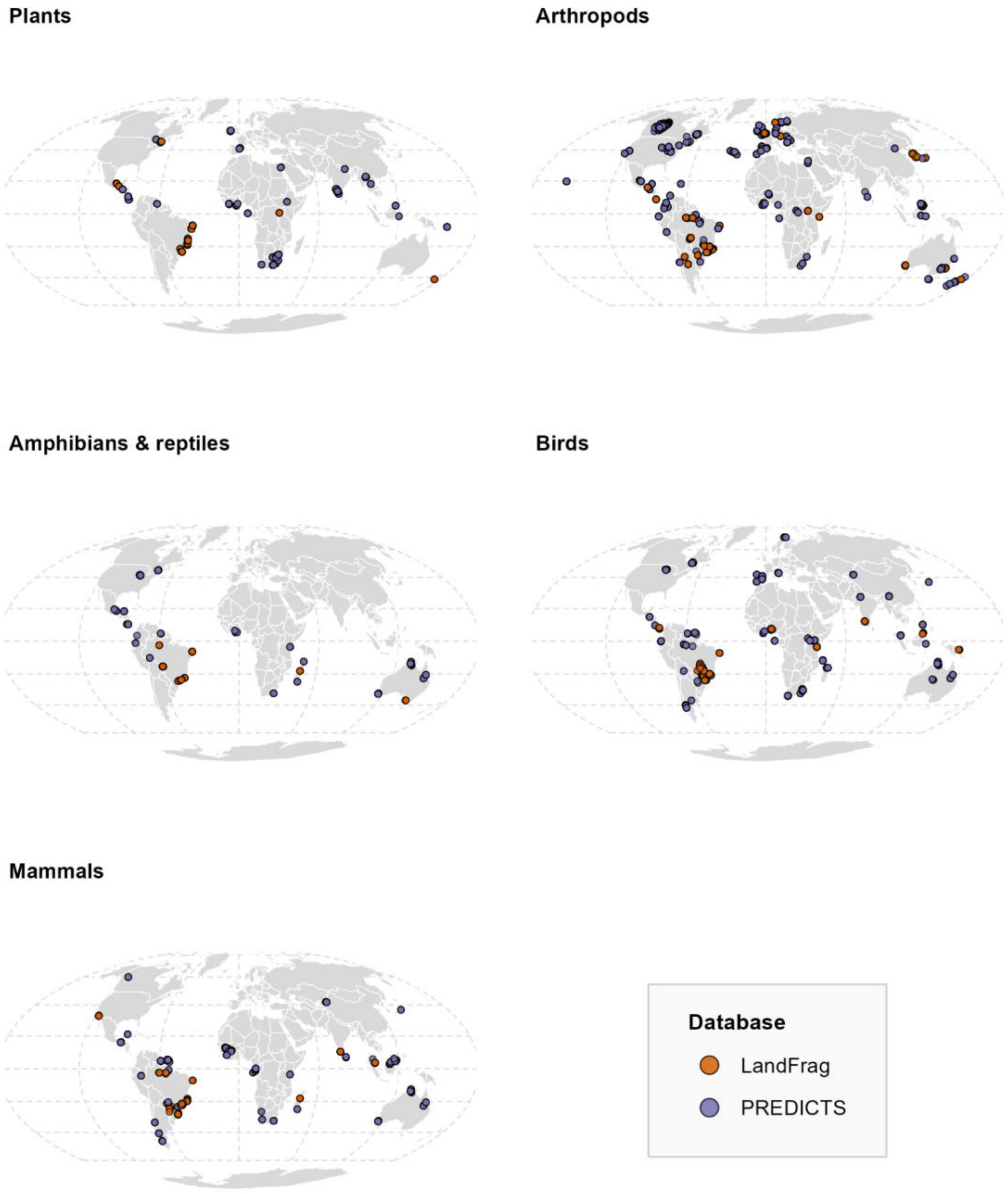


**Figure S1.** Global distribution of the sampling locations used to estimate local species richness for each taxonomic group. Panels show plants, arthropods, amphibians & reptiles, birds, and mammals. Each point represents the geographic centroid of a sampled continuous forest or forest fragment included in the analysis. Point color indicates the source database: LandFrag (orange) and PREDICTS (purple).

**Table S1**. Description of biodiversity metrics, ecosystem variables, and ecological-context predictors used in this study. Variables are grouped by analytical role, ecological dimension, scale, spatial resolution, and data source (see also figure 1, main text).

| Variable type | Metric | Ecological dimension | Analytical scale | Spatial resolution / unit | Source |
|---|---|---|---|---|---|
| Response | Local species richness | Biodiversity | Forest fragment / sampling location | Meters to few kilometers | LandFrag and PREDICTS |
| Response | Regional species richness | Biodiversity | Study | 1 × 1° | IUCN |
| Response | Ecosystem productivity (DHI-GPP) | Ecosystem functioning | Forest fragment | 1 km | DHI remote-sensing products |
| Predictor | Mean Species Abundance (MSA) | Biodiversity indicator | Forest fragment | 300 × 300 m | GLOBIO4 |
| Ecological context | NDVI | Vegetation condition | Forest fragment | 1 km | DHI remote-sensing products |
| Ecological context | EVI | Vegetation condition | Forest fragment | 1 km | DHI remote-sensing products |
| Ecological context | Forest cover | Landscape structure | Surrounding landscape | 1 km | Japan Aerospace Exploration Agency (FNF4) |
| Ecological context | Habitat proximity | Landscape structure | Surrounding landscape | 1 km | FNF4 |
| Ecological context | Biome | Bioclimatic context | Broad regional context | Biome polygon / categorical assignment | Olson *et al* 2021 |

**Table S2.** Overall relationships between Mean Species Abundance (MSA) and local and regional species richness, and ecosystem productivity (DHI-GPP). Estimates represent standardized MSA effects. For local richness and ecosystem productivity, coefficients represent within-study MSA effects, whereas the regional richness coefficient represents the within-biome MSA effect after accounting for broad differences among biomes and repeated studies. Within-study and within-biome fits are represented by the light grey lines in Fig. 2 (main text).

| **Ecological outcome** | ***β*** | **SE** | ***z*** | ***p*** |
|---|---|---|---|---|
| Local richness | 0.0354 | 0.00982 | 3.60 | <0.001 |
| Regional richness | −0.1166 | 0.0353 | −3.31 | <0.001 |
| Ecosystem productivity | 0.0250 | 0.00406 | 6.16 | <0.001 |

**Table S3.** Sensitivity analysis of MSA–local species richness relationships using abundance adjusted models. Taxon-specific changes in the effect of ecological context on the MSA–local species richness relationship are shown for the main models and models additionally accounting for within- and between-study variation in total abundance. Estimates represent the change in the MSA slope associated with a 1-SD increase in ecological context. Δ estimate is the abundance-adjusted estimate minus the main-model estimate.

| Context | Taxon | Main estimate [95% CI] | Main *p* value | Adjusted estimate [95% CI] | Adjusted *p* value | Δ estimate |
|---|---|---|---|---|---|---|
| NDVI | Plants | 0.152 [0.106, 0.198] | <0.001* | 0.095 [0.069, 0.121] | <0.001* | -0.057 |
| NDVI | Arthropods | 0.018 [-0.006, 0.042] | 0.143 | 0.004 [-0.013, 0.021] | 0.638 | -0.014 |
| NDVI | Amphibians & reptiles | -0.061 [-0.134, 0.012] | 0.103 | -0.027 [-0.085, 0.031] | 0.359 | 0.034 |
| NDVI | Birds | -0.036 [-0.065, -0.007] | 0.016* | -0.023 [-0.043, -0.003] | 0.023* | 0.013 |
| NDVI | Mammals | -0.000 [-0.043, 0.043] | 0.991 | 0.008 [-0.027, 0.044] | 0.64 | 0.009 |
| EVI | Plants | 0.099 [0.053, 0.144] | <0.001* | 0.084 [0.058, 0.109] | <0.001* | -0.015 |
| EVI | Arthropods | 0.019 [-0.003, 0.041] | 0.088 | -0.001 [-0.016, 0.014] | 0.909 | -0.02 |
| EVI | Amphibians & reptiles | 0.011 [-0.061, 0.084] | 0.764 | 0.005 [-0.050, 0.060] | 0.863 | -0.006 |
| EVI | Birds | -0.027 [-0.058, 0.004] | 0.093 | -0.010 [-0.031, 0.011] | 0.358 | 0.017 |
| EVI | Mammals | -0.021 [-0.065, 0.022] | 0.337 | -0.004 [-0.039, | 0.805 | 0.017 |

| | | | | | | |
|---|---|---|---|---|---|---|
| | | | | 0.031] | | |
| Forest cover | Plants | 0.017 [-0.025, 0.059] | 0.435 | 0.012 [-0.011, 0.035] | 0.32 | -0.005 |
| Forest cover | Arthropods | -0.002 [-0.029, 0.025] | 0.873 | -0.007 [-0.023, 0.009] | 0.412 | -0.005 |
| Forest cover | Amphibians & reptiles | -0.052 [-0.138, 0.034] | 0.238 | -0.018 [-0.087, 0.050] | 0.604 | 0.034 |
| Forest cover | Birds | 0.068 [0.033, 0.103] | <0.001* | 0.009 [-0.013, 0.032] | 0.412 | -0.058 |
| Forest cover | Mammals | -0.095 [-0.140, -0.049] | <0.001* | -0.033 [-0.069, 0.003] | 0.073 | 0.062 |
| Habitat proximity | Plants | -0.005 [-0.072, 0.063] | 0.895 | 0.029 [-0.009, 0.066] | 0.132 | 0.033 |
| Habitat proximity | Arthropods | -0.007 [-0.034, 0.020] | 0.587 | 0.000 [-0.013, 0.014] | 0.985 | 0.008 |
| Habitat proximity | Amphibians & reptiles | -0.031 [-0.158, 0.097] | 0.637 | -0.009 [-0.114, 0.095] | 0.86 | 0.021 |
| Habitat proximity | Birds | 0.006 [-0.043, 0.055] | 0.806 | -0.057 [-0.087, -0.028] | <0.001* | -0.064 |
| Habitat proximity | Mammals | -0.083 [-0.123, -0.042] | <0.001* | -0.030 [-0.060, -0.000] | 0.048* | 0.053 |

### References

Byrnes J E K and Dee L E 2025 Causal Inference With Observational Data and Unobserved Confounding Variables *Ecol. Lett.* **28** e70023

Chase J M, Blowes S A, Knight T M, Gerstner K and May F 2020 Ecosystem decay exacerbates biodiversity loss with habitat loss *Nature* **584** 238–43

Denomme-Brown S T, Cottenie K, Falls J B, Falls E A, Brooks R J and McAdam A G 2021 Examining the effects of heterospecific abundance on dispersal in forest small mammals *J. Mammal.* **102** 1484–96

Gonçalves-Souza T, Chase J M, Haddad N M, Vancine M H, Didham R K, Melo F L P, Aizen M A, Bernard E, Chiarello A G, Faria D, Gibb H, Lima M G, Magnago L F S, Mariano-Neto E, Nogueira A A, Nemésio A, Passamani M, Pinho B X, Rocha-Santos L, Rodrigues R C, Safar N V H, Santos B A, Soto-Werschitz A, Tabarelli M, Uehara-Prado M, Vasconcelos H L, Vieira S and Sanders N J 2025a Species turnover does not rescue biodiversity in fragmented landscapes *Nature* **640** 702–6

Gonçalves-Souza T, Vancine M H, Sanders N J, Haddad N M, Cortinhas L, Aase A L T O, de Aguiar W M, Aizen M A, Arroyo-Rodríguez V, Baz A, Benchimol M, Bernard E, Bertotto T J, Bispo A A, Bogoni J A, Boldorini G X, Bragagnolo C, Brosi B, Cantalice A S, do Carmo R F R, Cazeta E, Chiarello A G, de la Sancha N U, Didham R K, Faria D, Filgueiras B, Figueira J E C, Galvão G A, Garey M V, Gibb H, Gómez-Martínez C, González E, de Gusmão R A F, Henry M, de Jesus S, Kloss T G, Lázaro A, Leandro-Silva V, de Lima M G, da Silva Lima I, Lins-e-Silva A C B, Nally R M, Magalhães A R, Magnago L F S, Manu S, Mariano-Neto E, Mbora D N M, Melo F P L, Mutua M N, Neckel-Oliveira S, Nemésio A, Nogueira A A, Oliveira P M D A, Pádua D G, Paes L, de Paiva A B, Passamani M, Pena J C, Peres C A, Pinho B X, Pons J-M, Prasniewski V M, Reiniö J, dos Santos Rocha M, Rocha-Santos L, Rodal M J, Rodrigues R C, Safar N V H, Salomão R P, Santos B A, Santos M N, dos Santos J P, Savilaakso S, Schaefer C E G R, Silva M A M, da Silva F R, Silva R J, Simonelli M, Soto-Werschitz A, Stireman III J O, Storck-Tonon D, Szinwelski N, Tabarelli M, Teixeira C P, Totland Ø, Uehara-Prado M, Vaz-de-Mello F Z, Vasconcelos H L, Vieira S A and Chase J M 2025b LandFrag: A Dataset to Investigate the Effects of Forest Loss and Fragmentation on Biodiversity *Glob. Ecol. Biogeogr.* **34** e70015

Gotelli N J and Colwell R K 2001 Quantifying biodiversity: procedures and pitfalls in the measurement and comparison of species richness *Ecol. Lett.* **4** 379–91

Hudson L N, Newbold T, Contu S, Hill S L L, Lysenko I, De Palma A, Phillips H R P, Senior R A, Bennett D J, Booth H, Choimes A, Correia D L P, Day J, Echeverría-Londoño S, Garon M, Harrison M L K, Ingram D J, Jung M, Kemp V, Kirkpatrick L, Martin C D, Pan Y, White H J, Aben J, Abrahamczyk S, Adum G B, Aguilar-Barquero V, Aizen M A, Ancrenaz M, Arbeláez-Cortés E, Armbrecht I, Azhar B, Azpiroz A B, Baeten L, Báldi A, Banks J E, Barlow J, Batáry P, Bates A J, Bayne E M, Beja P, Berg

Å, Berry N J, Bicknell J E, Bihn J H, Böhning-Gaese K, Boekhout T, Boutin C, Bouyer J, Brearley F Q, Brito I, Brunet J, Buczkowski G, Buscardo E, Cabra-García J, Calviño-Cancela M, Cameron S A, Cancello E M, Carrijo T F, Carvalho A L, Castro H, Castro-Luna A A, Cerda R, Cerezo A, Chauvat M, Clarke F M, Cleary D F R, Connop S P, D'Aniello B, da Silva P G, Darvill B, Dauber J, Dejean A, Diekötter T, Dominguez-Haydar Y, Dormann C F, Dumont B, Dures S G, Dynesius M, Edenius L, Elek Z, Entling M H, Farwig N, Fayle T M, Felicioli A, Felton A M, Ficetola G F, Filgueiras B K C, Fonte S J, Fraser L H, Fukuda D, Furlani D, Ganzhorn J U, Garden J G, Gheler-Costa C, Giordani P, Giordano S, Gottschalk M S, et al 2014 The PREDICTS database: a global database of how local terrestrial biodiversity responds to human impacts *Ecol. Evol.* **4** 4701–35

Huete A, Didan K, Miura T, Rodriguez E P, Gao X and Ferreira L G 2002 Overview of the radiometric and biophysical performance of the MODIS vegetation indices *Remote Sens. Environ.* **83** 195–213

Liu D, Silveira E M O, Razenkova E, Anand A, Dubinin M, Hobi M, Pidgeon A M and Radeloff V C 2026 Global dynamic habitat indices (DHIs) based on MODIS and VIIRS vegetation products *Remote Sens. Environ.* **332** 115099

Nunez S, Arets E, Alkemade R, Verwer C and Leemans R 2019 Assessing the impacts of climate change on biodiversity: is below 2 °C enough? *Clim. Change* **154** 351–65

van de Pol M and Wright J 2009 A simple method for distinguishing within- versus between-subject effects using mixed models *Anim. Behav.* **77** 753–8

Radeloff V C, Dubinin M, Coops N C, Allen A M, Brooks T M, Clayton M K, Costa G C, Graham C H, Helmers D P, Ives A R, Kolesov D, Pidgeon A M, Rapacciuolo G, Razenkova E, Suttidate N, Young B E, Zhu L and Hobi M L 2019 The Dynamic Habitat Indices (DHIs) from MODIS and global biodiversity *Remote Sens. Environ.* **222** 204–14

Tomašových A, Kowalewski M, Nawrot R, Scarponi D and Zuschin M 2024 Abundance–diversity relationship as a unique signature of temporal scaling in the fossil record *Ecol. Lett.* **27** e14470

Xu Z, Li Y, Li B, Hao Z, Lin L, Hu X, Zhou X, Yu H, Xiang S, Pascal M-L-F, Shen W, He A, Chen L and Li Z 2023 A comparative study on the applicability and effectiveness of NSVI and NDVI for estimating fractional vegetation cover based on multi-source remote sensing image *Geocarto Int.* **38** 2184501
